# Sequencing-free whole genome spatial transcriptomics at molecular resolution in intact tissue

**DOI:** 10.1101/2025.03.06.641951

**Authors:** Yubao Cheng, Shengyuan Dang, Yuan Zhang, Yanbo Chen, Ruihuan Yu, Miao Liu, Shengyan Jin, Ailin Han, Samuel Katz, Siyuan Wang

**Author notes:** Department of Biological Sciences, Columbia University, New York, NY 10027, USA. These authors contributed equally to this work (Shengyuan Dang, Yuan Zhang).

## Abstract

Recent breakthroughs in spatial transcriptomics technologies have enhanced our understanding of diverse cellular identities, compositions, interactions, spatial organizations, and functions. Yet existing spatial transcriptomics tools are still limited in either transcriptomic coverage or spatial resolution. Leading spatial-capture or spatial-tagging transcriptomics techniques that rely on in-vitro sequencing offer whole-transcriptome coverage, in principle, but at the cost of lower spatial resolution compared to image-based techniques. In contrast, high-performance image-based spatial transcriptomics techniques, which rely on in situ hybridization or in situ sequencing, achieve single-molecule spatial resolution and retain sub-cellular morphologies, but are limited by probe libraries that target only a subset of the transcriptome, typically covering several hundred to a few thousand transcript species. Together, these limitations hinder unbiased, hypothesis-free transcriptomic analyses at high spatial resolution. Here we develop a new image-based spatial transcriptomics technology termed Reverse-padlock Amplicon Encoding FISH (RAEFISH) with whole-genome level coverage while retaining single-molecule spatial resolution in intact tissues. We demonstrate image-based spatial transcriptomics targeting 23,000 human transcript species or 22,000 mouse transcript species, including nearly the entire protein-coding transcriptome and several thousand long-noncoding RNAs, in single cells in cultures and in tissue sections. Our analyses reveal differential subcellular localizations of diverse transcripts, cell-type-specific and cell-type-invariant tissue zonation dependent transcriptome, and gene expression programs underlying preferential cell-cell interactions. Finally, we further develop our technology for direct spatial readout of gRNAs in an image-based high-content CRISPR screen. Overall, these developments provide the research community with a broadly applicable technology that enables high-coverage, high-resolution spatial profiling of both long and short, native and engineered RNA species in many biomedical contexts.

## Introduction

Recent inventions of spatial transcriptomics technologies have revolutionized biomedical research by enabling the profiling of gene activities across cells and tissues with spatial resolution^1-5^. Already widely adopted across various tissues, in many physiological and pathological contexts, these tools have provided unprecedented insights into cell identities, interactions, and tissue architecture, revealing key molecular mechanisms in health and disease^1-5^.

Existing methods can be broadly classified into: 1) spatial-capture/tagging technologies, which use high-throughput sequencing to read out spatially tagged transcripts; and 2) image-based technologies, which utilize multiplexed fluorescence *in situ* hybridization (FISH) or *in situ* sequencing (ISS) for direct spatial transcript detection^1-5^. Both families of approaches represent significant advancements, but are limited by trade-offs between transcriptomic coverage and spatial resolution: Spatial-capture/tagging techniques offer genome-wide transcript coverage but have lower spatial resolution and do not preserve cellular and sub-cellular morphologies^6-13^. In contrast, image-based methods provide single-molecule spatial resolution and maintain cellular and sub-cellular morphologies, but currently require pre-selection of a limited subset of transcripts for high-quality detection, restricting unbiased hypothesis generation^14-18^. Overcoming these major limitations is essential to allow for hypothesis-free and high-resolution spatial transcriptomics profiling.

While spatial transcriptomics can reveal molecular and cellular organization within tissues, it cannot determine direct functional mechanisms or causative factors. Combining these technologies with high-throughput CRISPR screens, which systematically perturb individual genes in single cells, offers a promising approach to identify functional regulators of spatial transcriptomic phenotypes. However, such advances require new methods for detecting *in situ* guide RNAs (gRNAs)—the synthetic RNA that targets specific genes for perturbation—to link perturbations to phenotypes.

Efforts to achieve more comprehensive and unbiased spatial transcriptome coverage within intact tissues have focused on improving ISS and multiplexed FISH methods compatible with intact tissue imaging. Key directions include developing more advanced probe design, signal amplification, barcoding, and imaging strategies to improve the accuracy, efficiency, and coverage of spatial transcriptomic profiling. For example, state-of-the-art targeted ISS techniques such as STARmap^17^ as well as some multiplexed FISH techniques (e.g. 10x Xenium) use padlock probes carrying barcodes to detect and amplify RNA signals through rolling circle amplification (RCA) for multiplexed visualization of transcript species in single cells. However, in these RCA-based methods the padlock oligo probes for templating the rolling circle reaction need to be individually synthesized and cannot be amplified *in vitro* for repeated use due to the design scheme^14,17-22^, limiting the scale of profiling to typically hundreds to ∼3000 genes^14,17-22^. Non-RCA methods RNA SeqFISH+ and MERFISH used amplifiable probe libraries and super-resolved image analysis or expansion microscopy respectively to achieve targeting of 10,000 RNA species, but were limited to detecting relatively long genes only, due to the “tiling” of each target transcript with tens of probes for enough signal-to-background ratio in single RNA detection^23,24^.

Here we introduce a method that achieves whole-genome-scale transcriptome coverage with single molecule spatial resolution, addressing these current issues of low spatial resolution or the need to pre-select a limited set of transcript targets. In addition to spatially profiling endogenous transcripts, our technology for the first time enables direct spatial readout of gRNA spacer sequences on individual gRNA molecules in pooled CRISPR screens, allowing high-performance mapping of genetic perturbations and their impact on cellular landscapes. These developments provide a broadly applicable tool to uncover the spatial transcriptome and its functions in diverse biomedical contexts.

## Results

### RAEFISH enables genome-wide spatial transcriptomic profiling at single-molecule resolution

To achieve genome-wide coverage in image-based spatial transcriptomics, we develop a multiplexed FISH method termed Reverse-padlock Amplicon Encoding FISH (RAEFISH) with several key innovations: First, most previous multiplexed FISH methods require tiling each RNA molecule with tens of primary FISH probes^15,16^, restricting the probing of short RNA species. High-performance ISS approach such as STARmap or RCA-based FISH methods (e.g. 10x Xenium) are less restrictive in terms of target RNA length, but use padlock probes with at least one of the two ends (5’ end) partially/completely hybridized to the target transcripts/cDNAs, thus requiring variable sequences at/close to the end^14,17-19^. This limits scaling up of the probe library due to cost (see details in the next paragraph). Unlike these methods, our design generates a “reversed” padlock – with the nick (two ends) of the padlock facing away from the transcript-interacting regions (allowing invariant ends), and with a priming probe (also with invariant ends) hybridized adjacent to it (Fig. 1A). This key element allows cost-efficient synthesis of probe libraries covering the entire transcriptome (see the next paragraph for details). Next, the two ends of padlock probes are ligated with the help of a splint oligo and DNA ligase (Fig. 1A). The splint oligo is removed by a toehold oligo after padlock ligation, and has an overhang sequence with special nucleotides at the 3’ end to prevent RCA priming from unremoved splint oligo in the next step. Next, RCA (depending on the specific hybridization of padlock and priming probes in adjacency) generates multiple copies of the target sequence originally bound by the padlock probe, and a library of encoding probes (also with invariant ends, thus compatible with cost-efficient synthesis) is hybridized to the amplified target sequences. Each encoding probe carries a combination of overhang sequences that uniquely encode the RNA identity (Fig. 1A). Finally, the combinations of encoding sequences are read out by sequential FISH, where dye-labeled readout probes are sequentially hybridized to the overhang sequences of encoding probes (Fig. 1A). The combinations of sequential readout signals produce “digital barcodes” that can identify the gene identities of the RNA molecules, as in RNA MERFISH^15^. The use of multiplexed FISH for decoding enables tunable signal density with a flexible coding scheme, minimizing signal overlap and improving decoding accuracy. For example, here by using a 94-choose-4 coding scheme (explained in detail below), on average only 4% (4/94) of target transcripts are imaged per imaging round. This low percentage helps reduce signal overlap and is critical for whole-transcriptome-scale profiling. In contrast, STARmap and other ISS methods image a high fraction (1/4) of target transcripts per round, which would cause significant signal overlap if probing the whole transcriptome^14,17,18,22^.

**Fig. 1.**
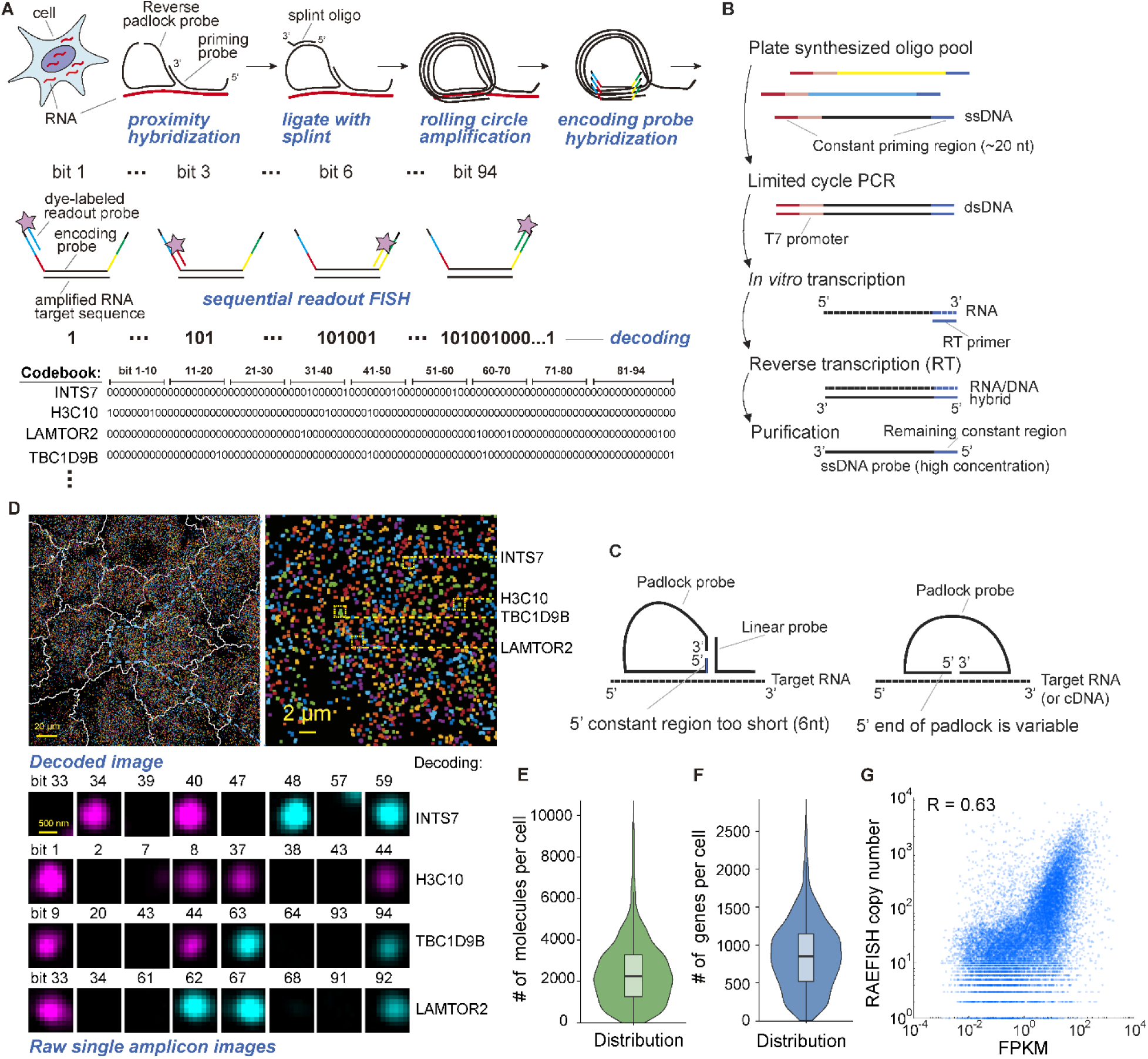
RAEFISH enables genome-wide spatial transcriptomic profiling at single-molecule resolution. **A**, Schematic illustration of the RAEFISH design. **B**, Schematic of oligo pool amplification. **C**, Demonstration of why other RCA-based ISS and multiplexed FISH methods are incompatible with the procedure in B. **D**, Example decoded image of A549 cells and raw single amplicon images from example molecules (yellow boxed in the zoom-in decoded image). Scale bars: 20 µm in decoded image, 2 µm in zoom-in decoded image, 500 nm in amplicon image. White lines in the decoded image indicate cell segmentation. The foci in the decoded images are pseudo-colored. The magenta and cyan colors in the raw single amplicon images correspond to Atto 647 and Alexa Fluor 750 fluorescent dyes respectively. **E-F**, Distribution of the numbers of RNA molecules (**E**) and different genes (**F**) detected in each cell. **G**, Correlation between RAEFISH and RNA-seq results.

To generate the priming, padlock, and encoding probe libraries in a cost-effective manner, we adapt an oligo pool amplification strategy previously developed for MERFISH encoding probe synthesis^15^ with minor modifications (Fig. 1B). In short, plate-synthesized template oligo pools undergo limited cycle PCR, *in vitro* transcription, reverse transcription, RNA degradation, and ssDNA probe purification. This approach allows a one-time purchase of plate-synthesized oligo pools, providing enough sequence-defined oligos to cover the entire transcriptome. These pools can be repeatedly amplified for multiple experiments, significantly lowering the probe cost per experiment. This procedure, however, leaves an ∼20-nt constant region at the 5’ end of the final probe (the reverse transcription priming region), and thus is incompatible with RCA padlock probe designs in ISS methods^14,17,18^, and in 10x Xenium^19^(Fig. 1C). As a result, current implementations of these other RCA-based methods use probes that are individually synthesized, which are cost-prohibitive to reach transcriptome scale. A RAEFISH probe library targeting 23,000 human genes (see results below) costed $5,132.40, which can support at least 2,000 experiments. Including the enzyme, reagent, and consumable cost of the probe amplification procedure, the per experiment cost is $158.18. In comparison, a STARmap probe library covering the same 23,000 genes would cost $58,220, supporting only 3 experiments without amplification. The per experiment cost would be $19,406.67 – 123-fold higher than RAEFISH.

Thus, RAEFISH combines the strengths of ISS and MERFISH to enable genome-wide targeting in imaged-based spatial transcriptomics: It creates an RCA-based design to allow 1) short transcript targeting, 2) cost-efficient probe synthesis, and 3) sequential FISH-based decoding to allow minimal signal overlap.

In a first test, we designed and synthesized a RAEFISH probe library targeting 23,312 human genes, including 16,501 protein-coding genes and 6,811 long-noncoding RNAs (Methods, Table S1). Each transcript was targeted by one pair of priming and padlock probes. Encoding probe design used a 94-choose-4 HD4 (Hamming distance 4) codebook, meaning each unique code consists of 94 binary digits, with exactly 4 set to “1” and the rest set to “0.”. The “1” digits indicate the specific rounds of readout imaging during which a targeted RNA species will be detected. This coding scheme also ensures that each pair of codes has a minimum Hamming distance of 4 (with at least 4-digit difference), allowing for the correction of any single-digit errors during the decoding process to improve detection. We tested this library and the RAEFISH procedure in fixed and permeabilized human lung adenocarcinoma A549 cells. Decoding involved 47 rounds of 2-color (647-nm and 750-nm laser excitable) readout FISH imaging (Fig. 1D), with fluorescent probes removed after each round using a 65% formamide wash. To correct for image drift, 0.1-µm green fluorescent beads were used as fiducial markers (Methods). On average, each cell had 2,443 decoded RNA molecules from 873 different genes detected (Fig. 1E-F), comparable to the detection efficiencies of single-cell RNA sequencing and high-quality ISS methods^17,18^. RNA copy numbers from RAEFISH correlated with FPKM values from bulk RNA-seq with a correlation coefficient of 0.63, validating the RAEFISH result (Fig. 1G).

### RAEFISH reveals cell-cycle associated genes and subcellular distributions of RNAs

To demonstrate the ability of RAEFISH to discover cell-cell heterogeneity in gene expression, we analyzed cell-cycle dependent gene expression changes in the proliferating A549 cell culture. Using an established cell cycle annotation pipeline for scRNA-seq analysis^25^, we called the cell-cycle phases (G1, S, G2/M) of individual A549 cells with the RAEFISH RNA copy numbers, and defined any expressed gene as a cell-cycle associated gene if its RNA copy number in one of the phases is significantly different from another phase. Based on this definition, our RAEFISH data showed 309 genes as significantly cell-cycle associated in proliferating A549 (Table S2). These genes could be hierarchically clustered into three expression patterns, representing upregulated marker genes in the G1, S and G2/M phases (Fig. 2A). Unsupervised clustering of single-cell RAEFISH RNA copy numbers of these genes yielded three clusters that corresponded to the cell cycle phases of the single cells (Fig. 2B). This list of cell-cycle associated genes includes known cell cycle markers, such as *CKS2*, *CENPF*, *CDC20* for the G2/M phase, and *RRM2*, *RRM1*, and *TYMS* for the S phase^26^ (Fig. 2A, Table S2). The list also includes genes that are less well known as cell cycle markers, but were reported to show cell-cycle dependent expression in certain contexts or control cell-cycle progression. For example, *PDE4A* which showed upregulation in G1 based on our data (Table S2) was reported to be selectively upregulated during G1 phase in C6 glioma cells^27^. *DUSP10*, *CHPF*, *SPRR2G*, and *ORAI3*, which showed upregulation during G1 in our data (Table S2), were all reported to control G1-to-S transition in different cell contexts^28-31^. Our data revealed multiple lncRNAs associated with different phases of cell cycle, including *LINC01270*, *ENSG00000260256*, *ENSG00000267372*, *LINC01719* upregulated in G1, *ENSG00000232358*, *RMDN2-AS1*, *SNHG17*, *ENSG00000273069*, *ENSG00000234296*, *TP53TG1*, *ENSG00000275672*, *LINC01788*, *ENSG00000267320*, *LINC01154*, *MIR4422HG*, *MILIP*, *ENSG00000280334* upregulated in G2/M, and *ENSG00000279179*, *ENSG00000258661*, and *ENSG00000278330* upregulated in S phase (Table S2). Among them, knockdown of *LINC01270* was reported to impair A549 growth^32^. Knockdown of *SNHG17* suppressed esophageal squamous cell carcinoma (ESCC) cell proliferation^33^. Knockdown of *TP53TG1* was shown to inhibit the proliferation of pancreatic ductal adenocarcinoma (PDAC) cells^34^. Knockdown of *MILIP* (c-Myc-Inducible Long noncoding RNA Inactivating P53) also reduced A549 growth^35^. As expected, gene ontology (GO) analysis of all the cell-cycle associated genes yielded top terms related to cell cycle, including e.g. leading strand elongation, deoxyribonucleoside biosynthetic process, DNA replication preinitiation complex assembly, positive regulation of chromosome condensation, metaphase/anaphase transition of cell cycle, etc (Fig. 2C).

**Fig. 2.**
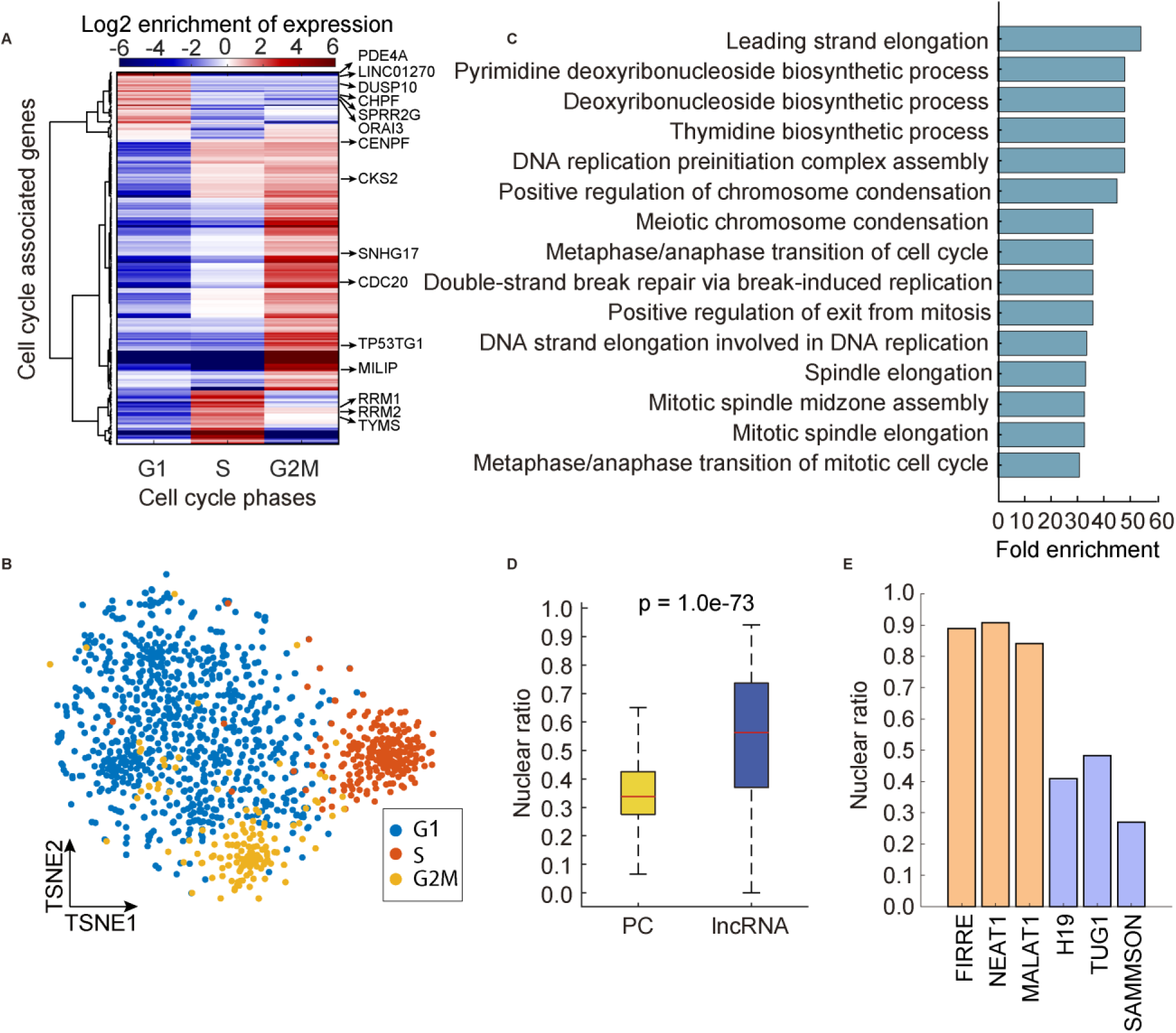
RAEFISH reveals cell-cycle associated genes and subcellular distributions of RNAs. **A**, Log2 enrichment of cell-cycle associated gene expression in G1, S, and G2M cells with hierarchical clustering. **B**, Unsupervised clustering with cell-cycle associated genes displayed with t-distributed stochastic neighbor embedding (TSNE). **C**. Enriched GO terms of all the cell-cycle associated genes. **D**. Nuclear ratios of protein coding (PC) RNAs and lncRNAs. **E**, Nuclear ratios of example lncRNAs. In all box plots throughout the manuscript, the boxes cover the 25th to 75th percentiles, the whiskers cover the 10^th^ to 90^th^ percentiles, and the line in the middle of the box represents the median value. P value in **D** was calculated by two-sided Wilcoxon rank sum test.

As a single-molecule method, RAEFISH allows sub-cellular spatial characterization of transcripts. To demonstrate this capability, we classified each detected copy of transcript as “nuclear” or “cytoplasmic” based on whether the x-y location of the transcript overlapped with nuclear DAPI staining, and calculated the nuclear ratio of each gene as the proportion of the transcripts being nuclear. As expected, lncRNAs on average showed higher nuclear ratio than protein coding genes (Fig. 2D). Classical nuclear lncRNAs e.g. *FIRRE*, *NEAT1*, and *MALAT1* showed high nuclear ratios, whereas lncRNAs known for cytoplasmic functions, e.g. *H19*, *TUG1*, and *SAMMSON*, showed lower nuclear ratios (Fig. 2E).

Overall, the analyses in this section demonstrate that RAEFISH uncovers meaningful cell-cell variations of gene expression associated with cell cycle and subcellular localizations of transcripts.

### RAEFISH uncovers spatial transcriptomic architectures and spatially dependent gene expression in liver tissue

To demonstrate the general utility of RAEFISH in mammalian tissue, we first adapted RAEFISH to adult mouse liver. We designed and synthesized a RAEFISH probe library targeting 21,955 genes in the mouse transcriptome, including 16,618 protein coding genes and 5,337 lncRNAs (Methods, Table S3). Fresh frozen tissue blocks were cryosectioned into 10-µm thick sections, formaldehyde-fixed, and permeabilized with Triton X-100. Then, the same RAEFISH procedure as in the cell culture experiment was applied. Raw RAEFISH images showed distinct amplicon fluorescent foci that were readily decodable (Fig. S1A). Decoded RAEFISH RNA copy numbers correlated with RNA-seq result at the bulk level, with a correlation coefficient of 0.74 (Fig. 3A), again validating the technique. Using both DAPI staining patterns and decoded RAEFISH images, we performed cell segmentation with a combination of the Cellpose^36-38^ and ClusterMap^39^ algorithms (Methods) (Fig. S1A). Segmented single cells contained on average 862 copies of detected RNA molecules per cell and 548 detected genes per cell (Fig. S1B-C). Again, these numbers are comparable to those obtained from some state-of-the-art sequencing-based single-cell/spatial transcriptomic technologies ^9,11,13,40,41^.

**Fig. 3.**
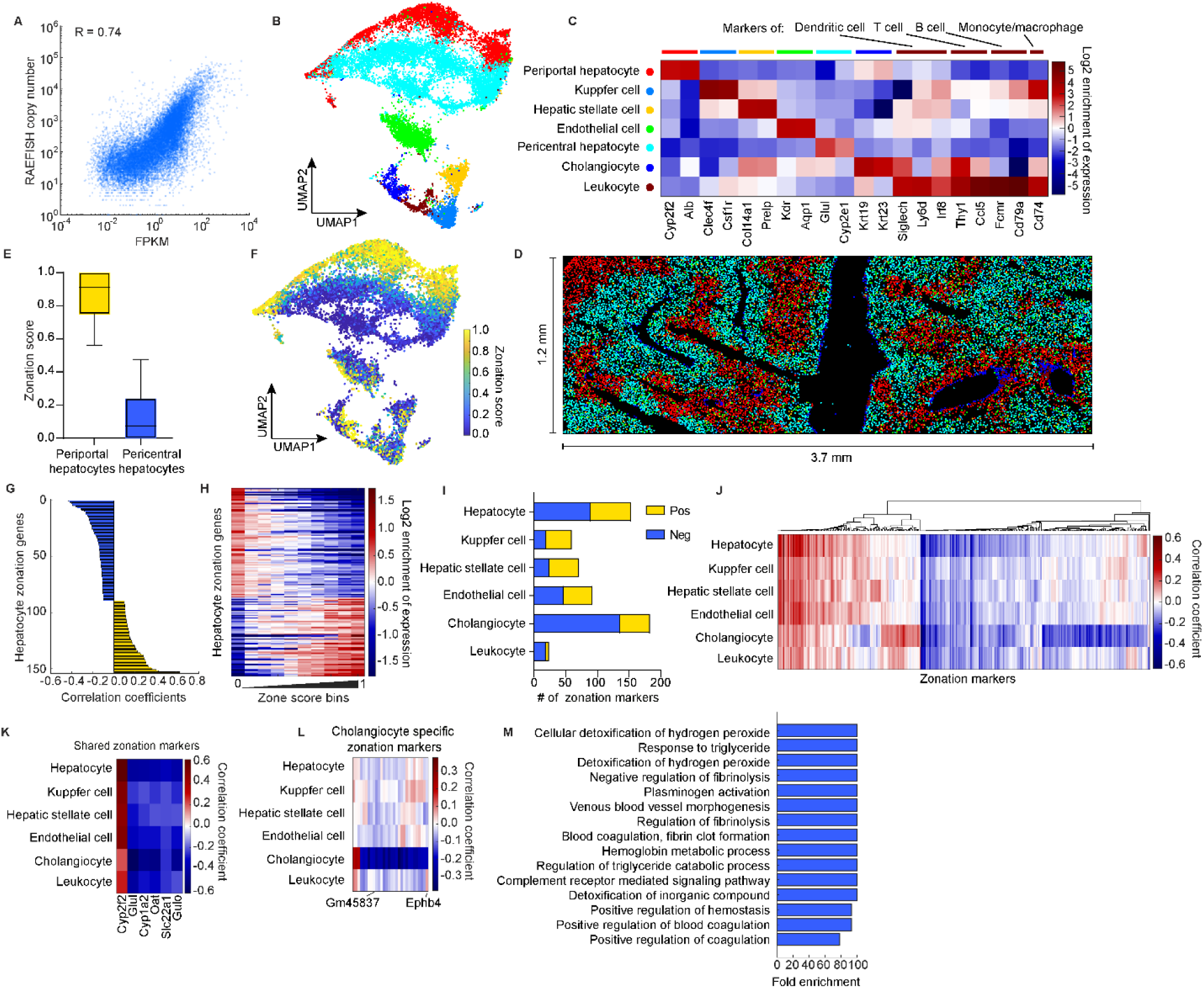
RAEFISH uncovers spatial transcriptomic architectures and spatially dependent gene expression in liver tissue. **A**, Correlation between RAEFISH and RNA-seq results. **B**, Single cell clusters displayed with Uniform Manifold Approximation and Projection (UMAP). **C**, Log2 enrichment of marker genes in each identified cell types. **D**, The *in-situ* map of the identified cell types. **E**, Zonation scores of periportal and pericentral hepatocytes. **F**, Zonation scores of all cells displayed on UMAP. **G**, Correlation coefficients between gene expression of hepatocyte zonation markers and zonation score. **H**, Log2 enrichment of expression of hepatocyte zonation markers in cells with binned zonation values. **I**, Number of zonation markers identified in each cell type. **J**, Expression-zone score correlation coefficients of zonation markers identified from all cell types with hierarchically clustering. **K**, Expression-zone score correlation coefficients of zonation markers shared by all cell types. **L**, Expression-zone score correlation coefficients of cholangiocyte specific zonation markers. **M**, Top GO terms of cholangiocyte-specific negative zonation markers.

We then computationally clustered the single cells in the RAEFISH dataset, plotted the clusters on a dimensionality-reduction UMAP plot, and annotated the cell type identities of each cluster. All major cell types in liver were successfully identified (Fig. 3B-C), with known marker genes expressed in the corresponding cell types (Fig. 3C). A well-known spatial feature of liver is liver zonation – hepatocytes (the major liver cell type) are spatially arranged into periportal (near portal veins) and pericentral (near central veins) zones^42-47^. Periportal and pericentral hepatocytes are known to differentially express zonation marker genes, in order to accommodate the different metabolic demands of the spatial zones^42,43,45-47^. Our single-cell transcriptomic clustering successfully identified periportal and pericentral hepatocytes (Fig. 3B), with the periportal hepatocytes marked by the expression of classical periportal zonation markers *Cyp2f2* and *Alb*^46,47^, and the pericentral hepatocytes upregulating classical pericentral markers *Glul* and *Cyp2e1*^46,47^ (Fig. 3C). When the cell identities were projected onto the tissue image, periportal and pericentral hepatocytes formed the expected, stereotypical zonation patterns (Fig. 3D). Note here the periportal and pericentral hepatocytes are transcriptionally defined, rather than conventionally defined by anatomy. As a result, a pericentral hepatocyte may not be closer to a central vein than to a portal vein or hepatic artery branch. Cholangiocytes, epithelial cells that line the bile ducts, also showed the expected lining pattern (Fig. 3D).

Given the gene expression differences between periportal and pericentral hepatocytes, we asked whether the other cell types also show distinct single-cell transcriptome profiles depending on the spatial zones they are located in. To associate each cell regardless of cell type with a spatial zone, we defined a zonation score for each cell as the percentage of nearby periportal hepatocytes among all nearby hepatocytes (Fig. S1D). Based on this definition, a cell with a higher zonation score (0.5-1) is more spatially associated with the periportal zone and a cell with a lower zonation score (0-0.5) is more associated with the pericentral zone. As a positive control, transcriptomically defined periportal hepatocytes showed higher zonation scores than pericentral hepatocytes (Fig. 3E). Projection of the single-cell zonation scores onto the single-cell transcriptome UMAP plot showed that the non-hepatocyte cell types also contained sub-clusters with different zonation scores (Fig. 3F), indicating distinct expression patterns associated with zonation. To identify in each cell type which genes are expressed in a zonation-dependent fashion, we calculated the correlation coefficient between the single-cell zonation scores and RAEFISH copy numbers of each gene (zonation correlation coefficient of each gene) among the cells of each type. The genes with significant correlations (positive or negative) were defined as zonation markers of that cell type (Fig. 3G). Positive zonation markers show higher expression in the periportal zone, while negative zonation markers are higher expressed in the pericentral zone (Fig. 3H). This analysis yielded 23-183 zonation markers for each cell type (Table S4), showing many genes’ expression profiles are zone-dependent and this phenomenon is not limited to hepatocytes. All major liver cell types have zonation markers (Fig. 3I).

To identify cell-type-specific and cell-type-independent zonation markers, we pooled the zonation markers identified from all cell types, and hierarchically clustered their zonation correlation coefficient profiles across the cell types. The result showed that hepatocyte zonation markers are often shared with other cell types (Fig. 3J). Several zonation markers are even shared by all major liver cell types (Fig. 3K). These cell-type-independent zonation markers include *Cyp2f2*, *Glul*, *Cyp1a2*, *Oat*, *Slc22a1*, and *Gulo*. On the contrary, some zonation markers are uniquely specific to one cell type (Fig. 3J). Particularly, cholangiocytes express many specific zonation markers not shared by other cell types (Fig. 3L). GO analysis showed that these cholangiocyte-specific zonation markers (particularly the 27 negative zonation markers associated with the pericentral zone) are enriched with top terms including cellular detoxification of inorganic compound, response to triglyceride, and blood coagulation – fibrin clot formation (Fig. 3M), corresponding to the pericentral zone’s functions in detoxification and lipogenesis and its association with ischemia-induced necrosis^42^.

We further analyzed finer spatial features in liver, focusing on cell-cell interactions. We asked which cell types are preferentially neighboring each other, and qualified the enrichment or depletion of the probability of each pair of cell types to be observed in a neighborhood of 20 µm radius, normalized by the probability expected from their abundance. As expected, periportal and pericentral hepatocytes showed depleted interactions, consistent with the zonation pattern (Fig. 4A). In addition, cholangiocytes and leukocytes preferentially interact with each other (Fig. 4A), consistent with previous reports of cholangiocyte-immune cell interaction^48^. To uncover potential gene expression changes in the interacting cholangiocytes and leukocytes that may underlie or result from the interactions, we first perform differential gene expression analysis comparing cholangiocytes neighboring leukocytes versus cholangiocytes not neighboring leukocytes (Fig. 4B, Table S5). Strikingly, all four upregulated genes in cholangiocytes neighboring leukocytes are components or chaperone of major histocompatibility complex (MHC) class II protein complex, including *H2-Eb1*, *H2-Aa*, *H2-Ab1*, and *Cd74* (Fig. 4B). This is consistent with reports that cholangiocytes can express MHC molecules and may perform antigen-presenting function^49^. Most of the down-regulated genes in cholangiocytes neighboring leukocytes are negative zonation markers in cholangiocytes (Table S5), e.g. *Ephb4*, *Lect2*, and *Gm45837* (Fig. 4B), indicating that the cholangiocyte-leukocyte interactions are preferentially located in the periportal region (and thus are depleted of pericentral markers). Indeed, when directly comparing the zonation scores of cholangiocytes interacting with leukocytes versus cholangiocytes not neighboring leukocytes, the former showed significantly higher zonation scores (Fig. 4C). Conversely, leukocytes interacting with cholangiocytes also showed higher zonation scores than leukocytes not interacting with cholangiocytes (Fig. 4D). We then analyzed the differentially expressed genes in leukocytes interacting with cholangiocytes compared to leukocytes not interacting with cholangiocytes (Fig. 4E, Table S6). GO analysis of top upregulated genes showed top enriched cellular component terms including elastic fiber, extracellular matrix, and extracellular space (Fig. 4F). Specifically, several top upregulated genes, e.g. *Fbln1*, *Mfap4*, *Ackr3*, and *Gpc6*, are components of extracellular matrix or cell surface receptors (Fig. 4E), which may mediate the interactions between cholangiocytes and these leukocytes. To identify which types of leukocytes preferentially interact with cholangiocytes, we analyzed the expression of leukocyte subtype marker genes in the interacting versus non-interacting leukocytes. We found macrophage markers *Ccl9*, *Cd14*, and neutrophil markers *Lcn2* and *Cd177* were preferentially expressed in the interacting leukocytes, whereas markers for T, B, NK, and dendritic cells were all depleted (Fig. 4G). These results suggest that macrophages and neutrophils expressing certain extracellular matrix proteins and surface receptors preferentially interact with MHC class II expressing cholangiocytes in the periportal zone of liver.

**Fig. 4.**
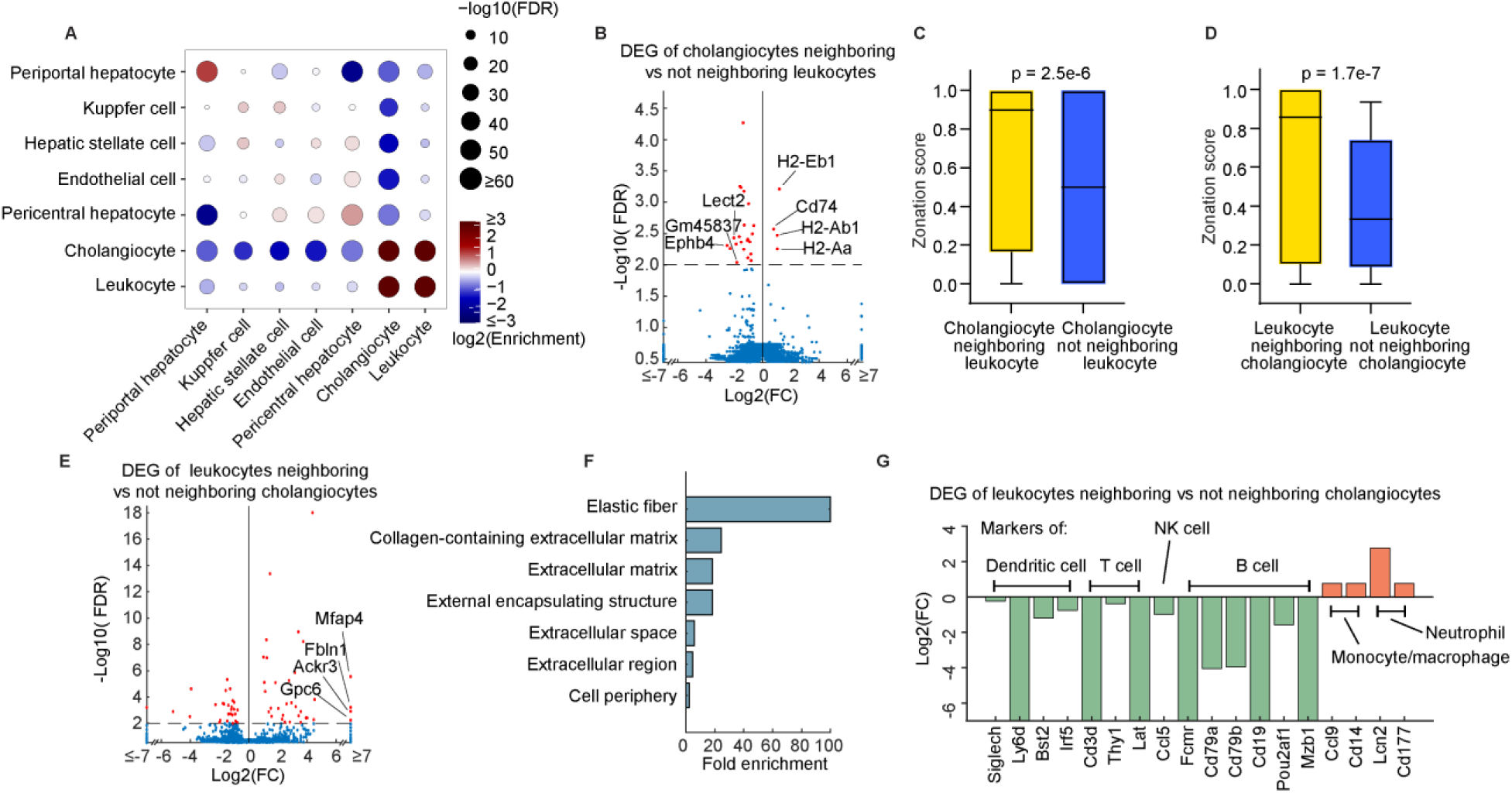
RAEFISH uncovers cell-cell communication in liver tissue. **A**, Log2 enrichment of cell-cell interactions. **B**, Differentially expressed genes (DEGs) of cholangiocytes neighboring leukocytes versus cholangiocytes not neighboring leukocytes. **C**, Zonation scores of cholangiocytes interacting with leukocytes versus cholangiocytes not neighboring leukocytes. **D**, Zonation scores of leukocytes interacting with cholangiocytes versus leukocytes not interacting with cholangiocytes. **E**, DEGs of leukocytes neighboring cholangiocytes versus leukocytes not neighboring cholangiocytes. **F**, Top GO terms of top upregulated genes in leukocytes neighboring cholangiocytes versus leukocytes not neighboring cholangiocytes. **G**, Log2 fold change of expression of leukocyte subtype marker genes in leukocytes interacting with cholangiocytes versus leukocytes not interacting with cholangiocytes. P value in **A** was calculated by Fisher’s exact test. P values in **C**, **D** were calculated by two-sided Wilcoxon rank sum test.

### RAEFISH uncovers characteristics of spatial transcriptomic architectures and cell-cell interaction in mouse placenta

To demonstrate that RAEFISH can be applied not only to mature organs but also during development, we applied the method to mouse placenta at embryonic day 13.5 (E13.5) using the same mouse RAEFISH probe library targeting 21,955 genes. We generated 10-µm thick longitudinal sections cryosectioned from fresh frozen placenta tissue block, followed by the same sample preparation protocol as for liver. The cross-section of half of a placenta was imaged, and the resulting RAEFISH raw image exhibited high quality RNA foci and were readily decodable (Fig. S2A). The decoded RAEFISH RNA copy numbers correlated with publicly available placenta bulk RNA-seq data^50^, with a correlation coefficient of 0.64 (Fig.S2B). After cell segmentation (Fig. S2A), we detected on average 1,266 RNA copies and 582 genes per cell (Fig. S2C-D).

Next, we used the single-cell RAEFISH RNA count profiles to identify cell type clusters and visualized the clusters on a UMAP dimensionality-reduction plot, annotated the cell type identity of each cluster based on known cell type marker genes^51-54^ (Fig. 5A-B), and then projected the cell type annotations back onto the spatial map of the placenta (Fig. 5C). Diverse cell types known to be present in placenta were identified (Fig. 5A-C). The identified cell types clearly delineate distinct zones in placenta, including (from top to bottom of the tissue image): the labyrinth, the junctional zone and the decidua^54,55^ (Fig. 5C). Decidual stromal cells (DSCs) and vascular endothelial cells (ECs) were identified in the decidua based on the RAEFISH results. In the junctional zone, spongiotrophoblasts (SPTs) were identified as the dominant cell type. Erythroblasts, sinusoidal trophoblast giant cells (S-TGCs), syncytiotrophoblast cells I (SynT-I), and syncytiotrophoblast cells II (SynT-II) were predominantly localized in the labyrinth (Fig. 5C). These spatial distribution and zonal organization of the cell types mapped by RAEFISH align with the expected organization as reported in literature^52,54,56^.

**Fig. 5.**
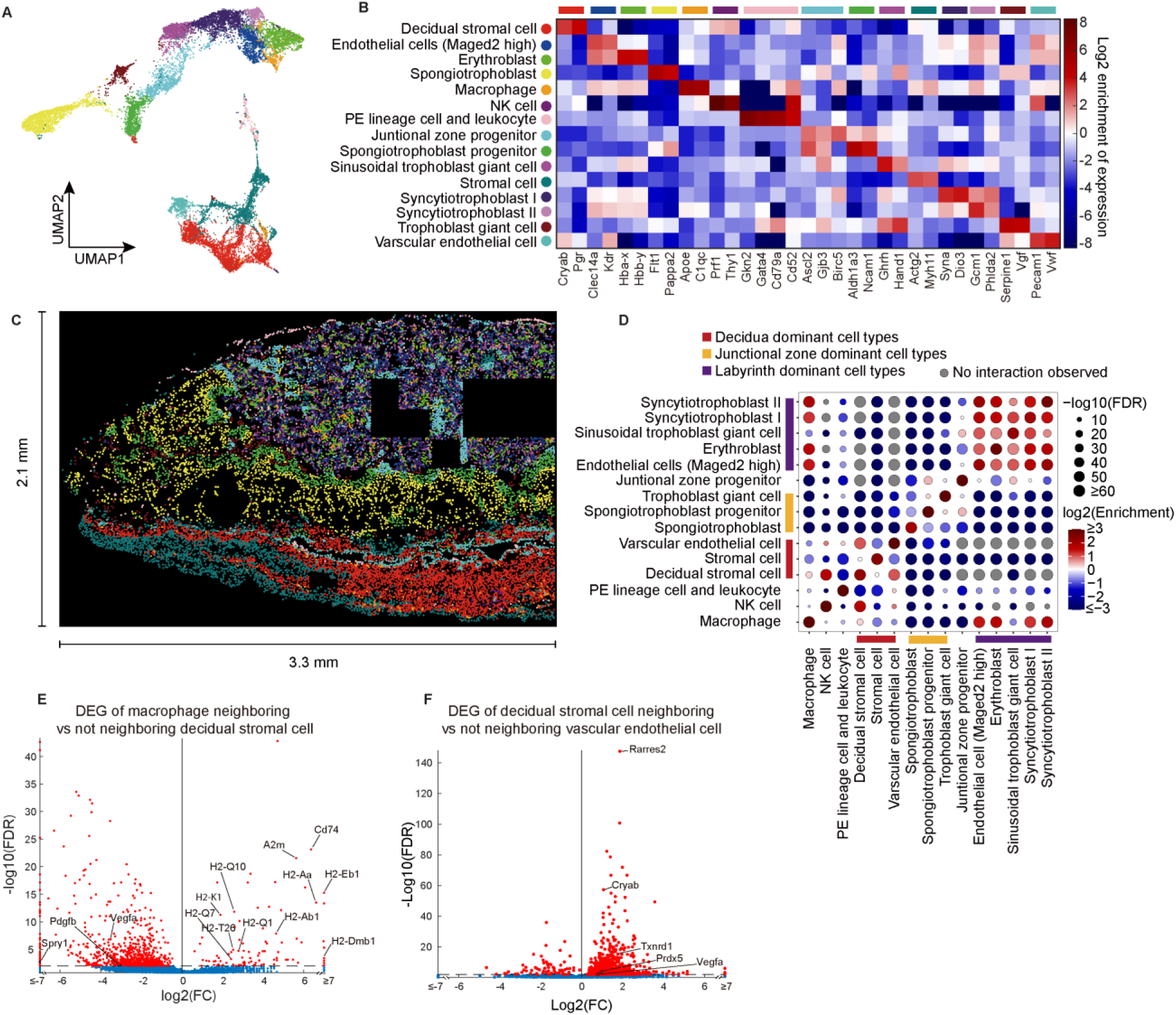
RAEFISH uncovers characteristics of spatial transcriptomic architectures and cell-cell interaction in mouse placenta. **A**, Single cell gene expression clusters displayed on UMAP. **B**, Log2 enrichment of marker genes in identified cell types. **C**, The *in-situ* map of the identified cell types. **D**, Log2 enrichment of cell-cell interactions. **E**, DEGs of macrophages neighboring decidual stromal cells versus macrophages not neighboring decidual stromal cells. **F**, DEGs of decidual stromal cells neighboring vascular stromal cells versus decidual stromal cells not neighboring vascular stromal cells. P value in **D** was calculated by Fisher’s exact test.

To further investigate finer spatial features in the mouse placenta, we conducted the cell-cell interaction analysis. As expected, cell types that were predominantly located in one zone showed depleted interactions with the cell types that were predominantly located in a different zone (Fig. 5D). We identified several enriched cell-cell interactions (Fig. 5D). For instance, in the decidua DSCs exhibited significantly enriched interactions with vascular ECs and natural killer (NK) cells, which corroborates previous publications^57,58^. Extensive cell-cell interactions were identified in the labyrinth, in line with the spatial map showing that these cells are highly intermingled in this region (Fig. 5C). In labyrinth, the interactions between ECs, erythroblast, SynT-I, SynT-II and macrophages are also consistent with their known functions and previous reports^58-62^.

To uncover the gene expression programs associated with the cell-cell interactions, we focused on several cell types with enriched interactions with DSCs (Fig. 5D). We first compared macrophages neighboring DSCs versus those that are not, as we observed a spatial segregation of macrophages into two groups, linked to their interaction status with DSCs: one group interacting with DSCs was located in the decidua, while the non-DSC-interacting macrophages were predominantly found in the labyrinth (Fig. S2E). Based on previous studies, macrophages that accumulate in the labyrinth (referred to as Hofbauer cells, HBCs) are fetal-derived macrophages, while maternal derived macrophages are primarily found in the decidua (referred to as decidual macrophages)^63,64^. We found that in comparison to HBCs, decidual (DSC-interacting) macrophages exhibited higher expression of MHC class II molecules and *Cd74* (Fig. 5E, Table S7), indicating a stronger pro-inflammatory phenotype^65^. This is consistent with previous reports showing that HBCs generally lack the M1 phenotype^66^. Intriguingly, we also found gene *A2m* highly expressed in decidual macrophages, which is a powerful anti-inflammatory molecule that can inhibit a variety of catabolic enzymes and cytokines^67,68^. These observations are consistent with previous reports that the same decidual macrophages can simultaneously express both pro-inflammatory and anti-inflammatory signatures^69^, related to their complex functions in placental development, including the control of maternal-fetal tolerance (anti-inflammatory) as well as trophoblast invasiveness (pro-inflammatory)^70-72^. In contrast, angiogenesis and vascular development related genes were upregulated in HBCs (Fig. 5E), such as *Vegfa*^73,74^ and *Pdgfb*^75^, consistent with prior knowledges that HBCs play an important role in labyrinth angiogenesis^61,63^. The Sprouty encoding gene *Spry1* was also higher expressed in HBCs (Fig. 5E). Sprouty has been reported as one of HBC’s secreted factors that modulates villous branching in human placenta^61,76^.

DSCs also exhibited significant interactions with vascular ECs, and differential gene expression analysis of DSCs adjacent to vascular ECs versus those not revealed genes associated with DSC functions (Fig. 5F, Table S8). For instance, *Rarres2* was upregulated in DSCs neighboring vascular ECs. This gene encodes the chemerin protein, a chemoattractant that can support peripheral blood NK cell migration through ECs and DSCs^77,78^. Consistently NK cells also showed enriched interactions with DSCs (Fig. 5D). *Vegfa* expression was also upregulated in DSCs neighboring vascular ECs, which is consistent with previous studies showing that VEGF is highly expressed from perivascular DSCs, and promotes EC proliferation and angiogenesis^79,80^. We also found three additional upregulated genes (*Cryab*, *Txnrd1* and *Prdx5*) in DSCs neighboring vascular ECs with known functions in protecting cells from oxidative stress induced by H_2_O_2_^81-83^ and protecting decidualization against stress conditions^84^. DSCs positioned near vascular ECs are exposed to oxidative stress products originating from maternal blood vessels^85^. Our results showed that vascular EC-interacting DSCs upregulate specific genes that help protect the cells against oxidative stress induced by H₂O₂.

In summary, here we demonstrated RAEFISH in mouse placenta, mapping sub-tissue zones, cell-cell interactions, and the interaction-associated gene expression in placenta. In the cell-cell interaction and the spatially dependent gene expression analysis, we linked the differential expression programs of macrophages in different zones to specific macrophage functions, and interpreted multiple gene expression signatures of DSCs neighboring ECs.

### RAEFISH uncovers spatial transcriptomic architectures in lymph node

To further demonstrate the general utility of REAFISH in different tissue contexts, we applied the 21,955-gene library to mouse lymph node, a more challenging tissue with small and crowded cells. To reduce the chance of overlapping cells in a tissue section, fresh frozen lymph nodes were cryosectioned into 8-µm thick sections and followed by the same sample preparation procedure as described above. We again obtained abundant and distinct RNA foci in this tissue (Fig. S3A). After decoding, the RNA copy numbers obtained from RAEFISH correlated with those obtained from scRNA-seq^86^ at the ensemble level (R = 0.65, Fig. S3B).

Next, we performed cell segmentation and clustered individual cells based on their RAEFISH count profiles. Major cell types in the mouse lymph node were identified as cell clusters and visualized on a UMAP plot (Fig. 6A-B). Each cell type showed strong expression of known marker genes (Fig. 6B). Two groups of B cells were identified, both exhibiting strong expression of classic B cell markers but differing in their cellular locations (Fig. S3C-D). They were assigned as marginal zone B cells and follicular B cells^87^. The most obvious spatial feature of the lymph node is the separation of B cell and T cell zones. B cells are mainly located in the cortex, the outer region of the lymph node and T cells are mainly located in the paracortex, the inner region of the lymph node^88^. By projecting the cell identities onto the tissue map, we see a clear spatial separation of B cell and T cell zones as expected (Fig. 6C). B cell zones are known to contain specialized microstructures known as germinal centers (GCs), where B cells undergo proliferation and somatic hypermutation^89^. On our tissue map, the oval-shaped GC B cell regions are clearly visible (Fig. 6C). GC B cells exhibited expression of cell proliferation markers, including *Mki67*, *Aurkb*, and *Top2a*, as well as *Aicda*, the gene encoding activation-induced cytidine deaminase (AID), which is critical for somatic hypermutation (Fig. 6B). Two distinct groups of plasma cells were further identified with distinct gene expression specialized for IgA and IgG antibody production (Fig. 6B). The T cell zone, located in the inner region of the lymph node, was enriched with dendritic cells, consistent with prior knowledge that dendritic cells are primarily located in the paracortex of lymph nodes^90^ (Fig. 6C, S3E).

**Fig. 6.**
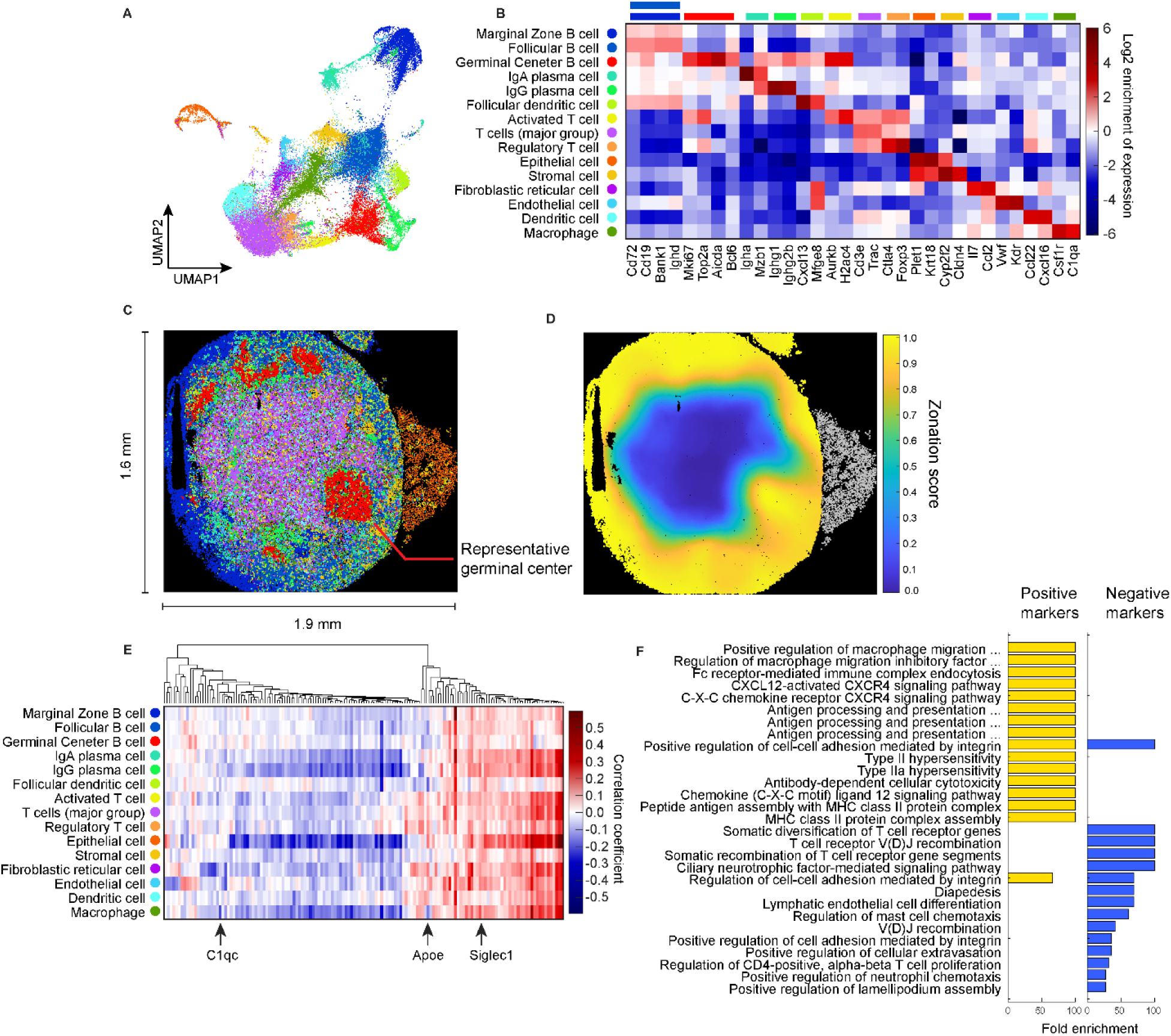
RAEFISH uncovers spatial transcriptomic architectures in mouse lymph node. **A**, Single cell gene expression clusters displayed on UMAP. **B**, Log2 enrichment of marker genes in each identified cell type. **C**, The *in-situ* map of the identified cell types. **D**, Zonation scores of the cells in the lymph node. Cells in gray colors were not involved in zonation analysis. **E**, Expression-zone score correlation coefficients of all zonation markers with hierarchically clustering. **F**, Top GO terms of positive and negative zonation markers.

To further analyze the spatial distribution of identified cell types and the spatial dependence of their gene expression, we defined a lymph node zonation score for each cell similar to the liver zone score definition (Fig. S1D). This score represents the percentage of nearby B cells relative to the total number of nearby B cells and T cells. As expected, we see a clear separation of B cell and T cell zones by projecting the zone score onto the tissue map (Fig. 6D). According to this definition, a cell with a higher zonation score (0.5–1) is more spatially associated with the B cell zone, while a cell with a lower zonation score (0–0.5) is more associated with the T cell zone. To analyze the zonation-dependent gene expression, we calculated the correlation between gene expression with the zone score in each cell type. The genes with significant correlations (positive or negative) from each cell type were grouped and plotted (Fig. 6E). Hierarchical clustering showed two major group of zonation markers. Gene Ontology analysis showed that these genes are closely related with immune regulation and cell differentiation in lymph nodes (Fig. 6F). The positive zonation markers are related with CXCL12-CXCR4 signaling pathway, which is crucial for germinal center organization^91^. The negative zonation markers are related with T cell differentiation, such as T cell receptor recombination (Fig. 6F). In addition, the zonation markers identified in specific cell types provide insights into their differential functions across distinct lymph node regions. For instance, in macrophages, we observed a significant negative correlation between the expression of *C1qc* and *Apoe* and the zone score, indicating that these genes are preferentially expressed in the T cell zones (Fig. 6E). In contrast, *Siglec1* in macrophages exhibited a significant positive correlation with the zone score, suggesting its preferential expression in B cell zones (Fig. 6E). These observations align with previous studies, which report that *Apoe* and *C1qc* are highly expressed in macrophages residing in the T cell zone, where they play roles in regulating T cell function^92,93^, whereas *Siglec1*+ subcapsular sinus macrophages are located at the outer edge of the lymph node and interact with lymph-borne antigens^94-96^. *Siglec1* encodes sialoadhesin, which can function as an endocytic receptor that mediates macrophage-pathogen adhesion and facilitates phagocytosis^97^.

Overall, our application of RAEFISH in three distinct tissue types demonstrates the general utility of this method to uncover spatial features in mammalian tissue at the levels of both larger sub-organ zones and finer cell-cell interactions and to discover underlying gene expression features associated with the spatial organization and cell-cell communication in a largely unbiased fashion.

### RAEFISH enables direct readout of gRNAs in an image-based CRISPR screen

Spatial omics technologies have significantly advanced our understanding of molecular and cellular organizations within tissues, but they fall short in revealing direct functional mechanisms or causative factors. In contrast, high-throughput genetic screens, which can systematically perturb numerous individual genes at once in single cells, offer a powerful approach to uncovering genetic regulators driving specific phenotypes. Thus, there is a critical need to develop high-throughput, high-content functional genomics screens with spatial omics readouts. Several methods combine lentiviral pooled CRISPR screen with spatial omics by pairing a barcode with each gRNA and reading out of the barcode with ISS or multiplexed FISH for gRNA identification^98-102^. However, these methods may suffer from RNA genome recombination during lentiviral packaging^103^, leading to shuffled gRNA-barcode pairings that reduce screening sensitivity and accuracy^103^. Some pipelines are also experimentally complex, requiring additional *in vitro* sequencing and integration of multiple barcode RNA molecules per cell for full decoding^99,101^. Recent works used T7 *in situ* transcription to amplify gRNA sequences from its integrated genomic locus for ISS or multiplexed FISH detection, avoiding the need of paired barcodes^104,105^. However, this generates only one gRNA imaging spot per cell nucleus from its DNA locus, instead of directly imaging the *in vivo* expressed gRNAs. Here we develop Perturb-RAEFISH to directly read out gRNAs from the numerous gRNA copies per cell in image-based high-content CRISPR screens.

We first established a pooled CRISPR knockout cell library with A549 cells, individually knocking out 574 expressed DNA/chromatin-associated-protein-coding genes in this cell line, with 3 gRNAs for each target gene and additional nontargeting/intergenic control gRNAs (Methods, Table S9). To test the direct RAEFISH readout of numerous gRNAs in this cell library, we designed RAEFISH padlock and linear probes targeting the gRNA spacer (the sequence that defines the genomic target to be modified) and scaffold regions (Fig. 7A). After splint-oligo-mediated ligation of the padlock probe and RCA, the gRNA spacer sequence is significantly amplified for multiplexed FISH encoding and decoding, as in native RNA RAEFISH (Fig. 1A). To assign detected gRNAs to individual cells, and in turn identify perturbations in the cells, we imaged whole cells with total protein stain, and segmented the cell bodies (Methods). Importantly, our cell library was constructed with the CROP-seq strategy that led to gRNA expression from both an RNA pol II promoter and a pol III promoter, resulting in many copies of gRNAs in each cell’s nucleus and cytoplasm^106^ (Fig. 7B). We achieved the detection of 41 copies of gRNAs per cell on average, and 89.3% of cells had one dominant gRNA identity (Fig. 7C-D).

**Fig. 7.**
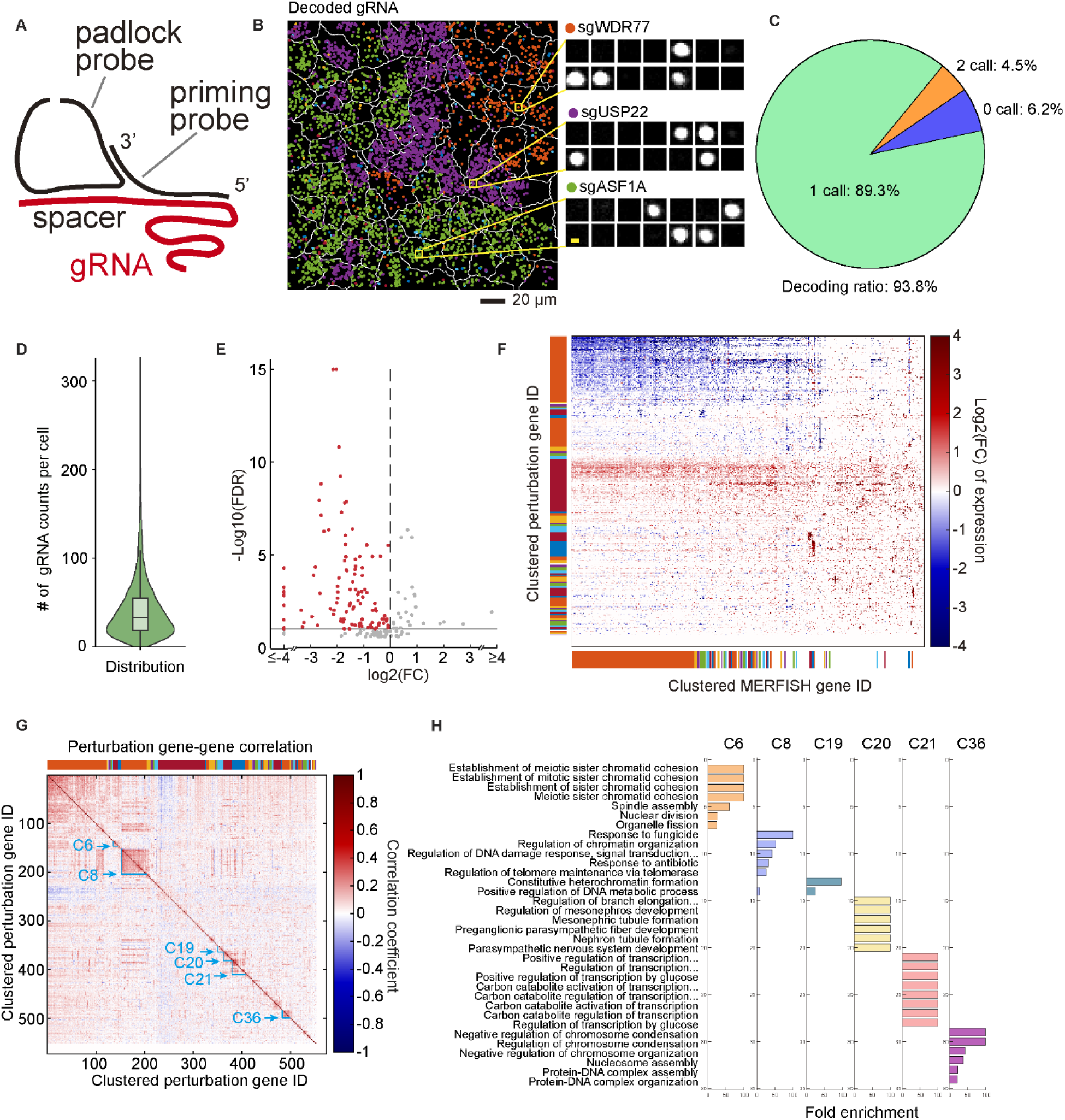
RAEFISH enables direct readout of gRNAs in an image-based CRISPR screen. **A**, Schematic illustration of direct RAEFISH targeting of gRNA spacer region. **B**, Representative decoded gRNA molecules in single cells. Different pseudo colors represent different gRNA identities. White lines show cell segmentation. Each gRNA is encoded with a 14-bit binary code with 4 “1” bits. **C**. Pie chart representing the percentages of cells with different numbers of dominant gRNA identities. **D**, Distribution of the copy numbers of gRNAs detected in each cell. **E**, Log2FC of expression (from RNA MERFISH counts) of perturbation target genes (comparing cells with perturbation gRNAs versus cells with negative control gRNAs). **F**, Log2FC of all gene expression detected by RNA MERFISH in all detected perturbations. Both the perturbation target genes and MERFISH probed genes are hierarchically clustered (color bars along y and x axes). **G**, Correlation coefficients of MERFISH gene expression profiles between each pair of perturbations. Six representative perturbation gene clusters are indicated by arrows. **H**, Top GO terms of the highlighted clusters in G.

We further used RNA MERFISH to simultaneously detect endogenous RNAs. For RNA MERFISH we designed and synthesized probes targeting 492 RNAs encoding DNA/chromatin-associate proteins, including 183 of the CRISPR knockout targets. Application of the RNA MERFISH probe set in combination with gRNA readout by RAEFISH validated reduction of most of the probed knockout target RNAs, likely through nonsense mediated decay (Fig. 7E).

By comparing with the negative controls, we found 552 of the perturbation target genes significantly (FDR<0.05) affected the expression of one or more MERFISH-probed genes (Fig. 7F). Clustering of perturbed transcriptome profiles revealed 52 clusters of the perturbation target genes (where each cluster contains target genes that generated similar perturbation effects) and 43 clusters of the MERFISH-probed genes (where each cluster contains probed genes with similar responses to the perturbations) (Fig. 7F, Table S10). Focusing on the perturbed genes read out by RAEFISH, a gene-gene correlation analysis showed that genes in the same clusters were more correlated in their knockout response (Fig. 7G). GO analyses showed that the clusters were associated with specific biological processes (Fig. 7H). For example, Cluster 6 is associated with establishment of sister chromatid cohesion, spindle assembly and nuclear division. Cluster 8 is associated with DNA damage response and regulation of telomere maintenance via telomerase. Cluster 19 is associated with constitutive heterochromatin formation. Cluster 20 is associated with nephron tubule formation and parasympathetic nervous system development. Cluster 21 is associated with regulation of transcription from RNA polymerase II promoter by glucose. And Cluster 36 is associated with regulation of chromosome condensation and nucleosome assembly (Fig. 7H).

Overall, our results here demonstrate the versatility of RAEFISH in detecting short, engineered RNAs in image-based, high-content genetic screening.

## Discussion

In summary, RAEFISH provides researchers with the capacity to profile spatial transcriptome with genome-scale coverage and single-molecule spatial resolution in cells and intact tissues. This first demonstration of the technology in various cell and tissue contexts yielded a list of biological insights. Particularly, the human cell culture RAEFISH experiment discovered dozens of unannotated/under-annotated long noncoding RNAs that are associated with cell cycle, and revealed subcellular localizations of transcripts. The mouse liver RAEFISH identified cell-type-specific and cell-type-invariant zonation markers across all major liver cell types, and uncovered a periportal-zone-enriched interaction between cholangiocytes and immune cells that likely supports an antigen presenting function of cholangiocytes. The mouse placenta RAEFISH highlighted the different characteristics of macrophages in labyrinth and decidua, and reveals the functional significance of cell-cell interaction between DSCs and vascular ECs. The mouse lymph node RAEFISH revealed spatial gene expression profiles associated with the B and T cell zones and the GC structure. Finally, besides the endogenous transcriptome, we showed that the RAEFISH technology is also applicable to engineered RNAs, enabling direct readout of gRNAs in an image-based Perturb-RAEFISH CRISPR screen, which solves a key limitation in the field. To our best knowledge, this work represents the first time that more than 20,000 RNA species were directly probed and imaged *in situ* with any technology, and the first time numerous different gRNAs were directly probed and distinguished by imaging with single molecule resolution in cells.

As the first introduction of the RAEFISH technology, a limitation of this work is we have not incorporated all approaches and modalities available in the rapidly developing spatial transcriptomics field. For example, RCA in combination with tissue clearing has been shown to enable thick-tissue spatial transcriptomic imaging^17,107^. As RAEFISH already includes the RCA signal amplification component, we expect further combining RAEFISH with tissue clearing will enable whole-genome-scale spatial transcriptomic imaging in thick tissue. Furthermore, expansion microscopy was shown to improve the quality of image-based spatial transcriptomics^18,24^. We expect RAEFISH to be fully compatible with expansion microscopy and such a combination may further improve the transcriptomic coverage of the technique to e.g. short noncoding RNA varieties. For RNA species that are too short to support the binding of even one pair of reverse padlock and linear priming probes, a recent technique to extend the length of such RNAs through *in situ* polyadenylation may help^108,109^. Finally, while we only demonstrated Perturb-RAEFISH in cell culture, the technique should be adaptable to *in vivo* screens, given recent *in vivo* demonstrations of RCA and FISH-based readout of barcode mRNAs paired with gRNAs^102,110^.

In sum, the development of RAEFISH provides the biomedical research community with a generalizable research tool, which will help bring more spatial and mechanistic insights across health and disease.

## Acknowledgments

We thank Xinlin Zeng and Eva Wang for help with experiments, members of the Wang lab for helpful discussions. We thank members of the Yale Center for Genome Analysis and the Keck DNA Sequencing facility for their help. Figures S1D were created in part in BioRender.com. S. K. was in part supported by NIH (R21CA282629). S. W. was partly supported by NIH (UH3CA268202, U01CA260701, R01HG011245, DP2GM137414, R01CA292936, R01HG012969, R01HG013503, P50CA196530-10S1), Pershing Square Sohn Cancer Research Alliance, American Federation for Aging Research and Hevolution Foundation. This work was in part supported by NIH (DP2GM137414, R01HG011245, R01HG013503, UH3CA268202) and Pershing Square Sohn Cancer Research Alliance.

## Author contributions

S.W. conceived the study. S.W. and Y. Cheng designed the experiments. Y.Cheng., S.D., Y.Z., Y.Chen., R.Y., M.L., S.J., A.H., S.K., and S.W. performed experiments. Y.Cheng., S.D., Y.Z., and S.W. analyzed data. Y.Cheng., S.D., and S.W. wrote the manuscript with comments from all authors.

## Declaration of Interests

S.W. and Y.Cheng are inventors on a patent application filed by Yale University related to RAEFISH and Perturb-RAEFISH reported in this manuscript. S.W. is one of the inventors on a patent applied for by Harvard University related to MERFISH.

---

**Fig S1.**
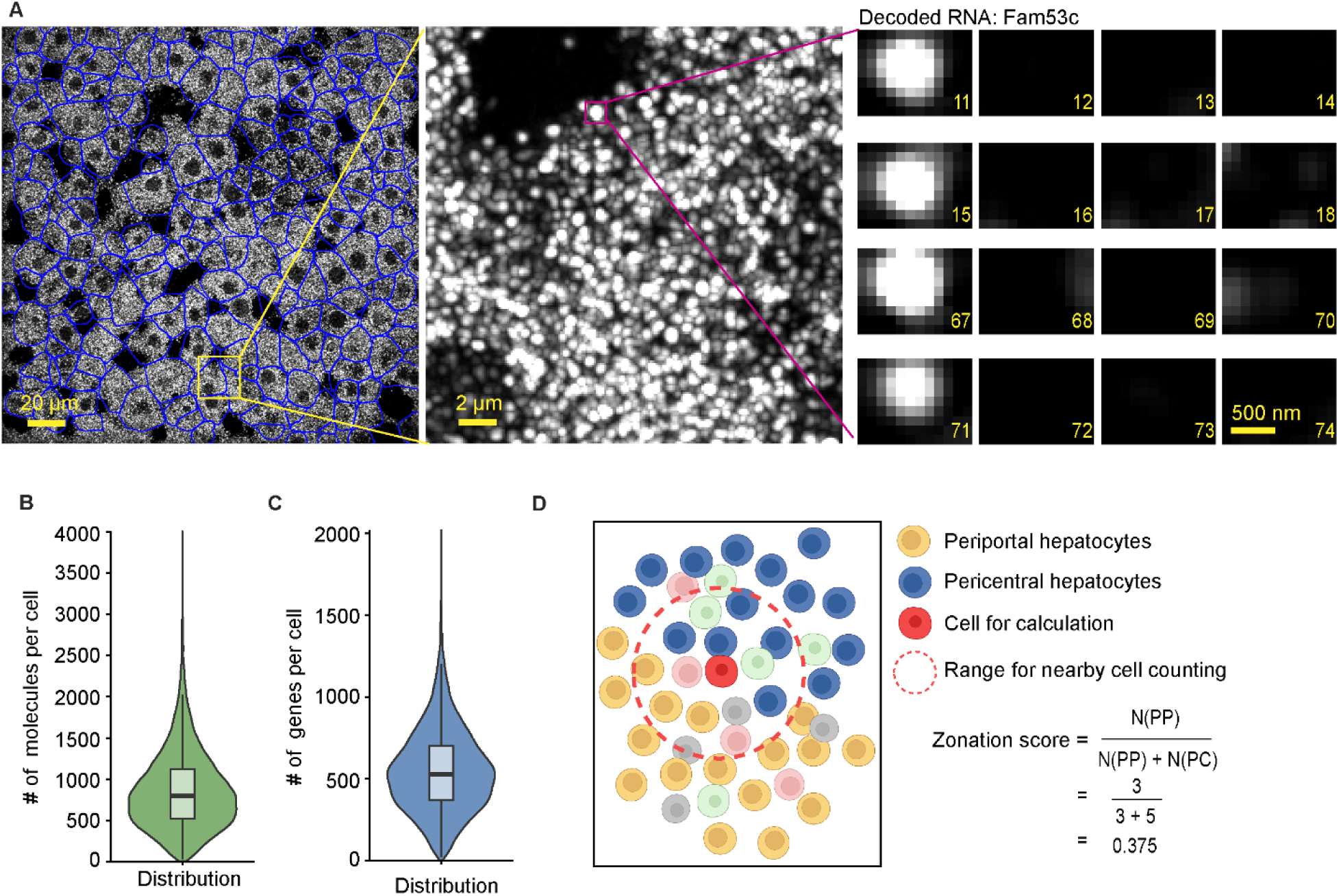
RAEFISH in mouse liver tissue. **A**, An example of raw RAEFISH foci and decoding result in mouse liver. Blue lines indicate cell segmentation result. Scale bars: 20 µm in the left image, 2 µm in the middle zoom-in image, 500 nm in the right amplicon images. **B-C**, Distribution of the number of RNA molecules (**B**) and different genes (**C**) detected in each cell. **D**, Schematic illustration of zonation score calculation.

**Fig S2.**
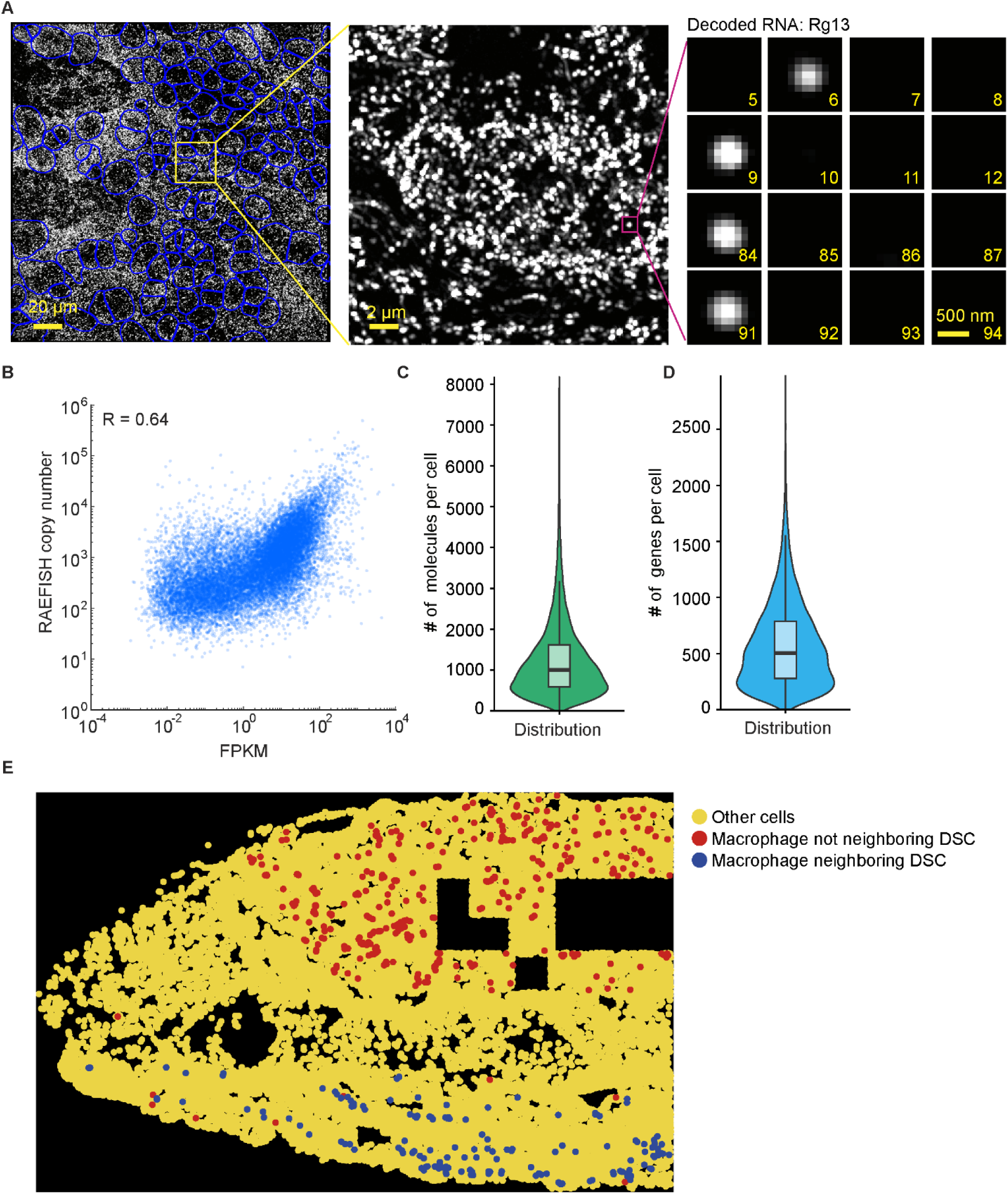
RAEFISH in mouse placenta. **A**, An example of raw RAEFISH foci and decoding result in mouse placenta. Blue lines indicate nuclear segmentation result. Scale bars: 20 µm in the left image, 2 µm in the middle zoom-in image, 500 nm in the right amplicon images. **B**, Correlation between RAEFISH and RNA-seq results. **C-D**, Distribution of the number of RNA molecules (**C**) and different genes (**D**) detected in each cell. **E**, Spatial distribution of detected macrophages that were neighboring DSCs (blue) or not (red).

**Fig. S3.**
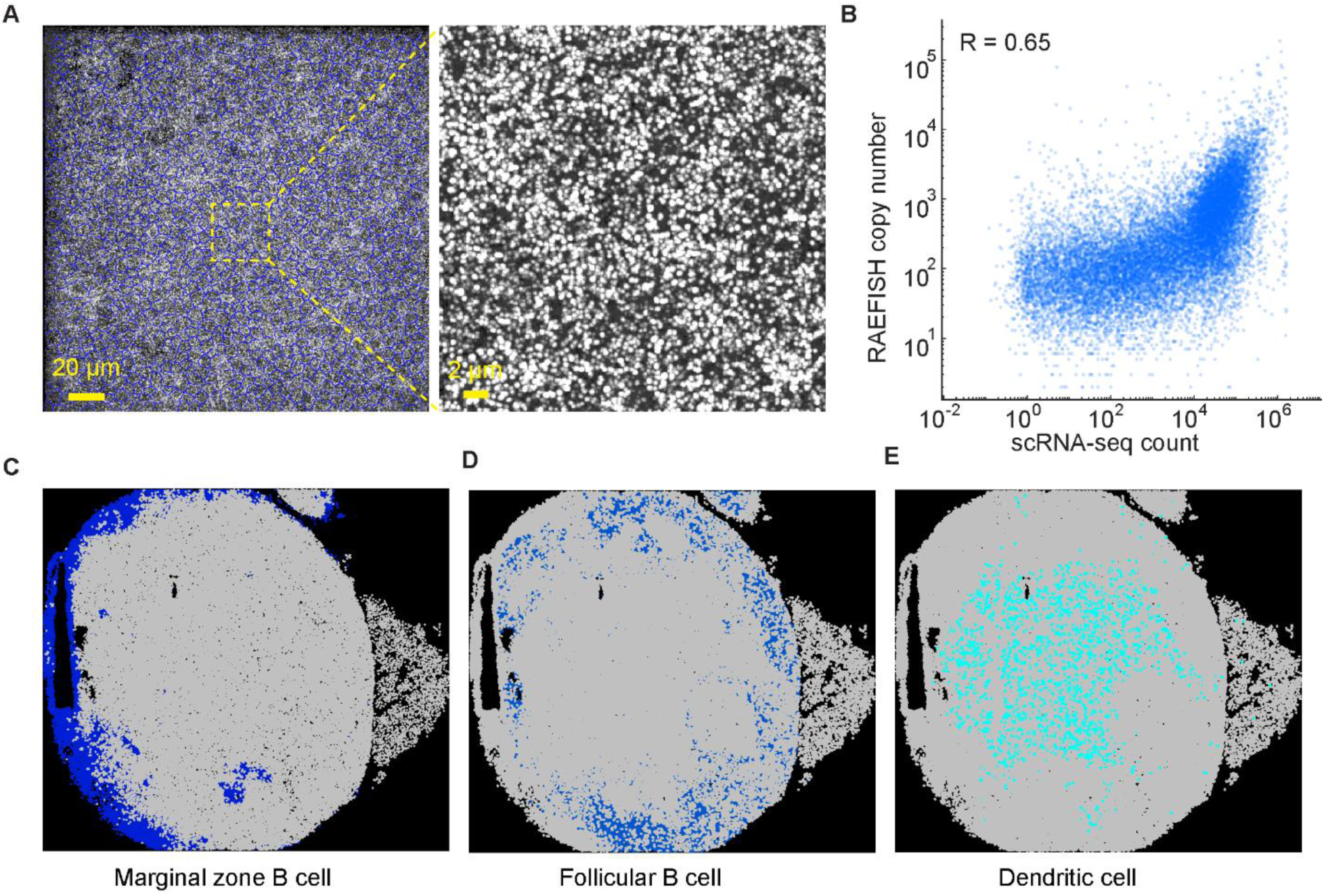
RAEFISH in mouse lymph node tissue. **A**, Example raw RAEFISH foci in mouse lymph node. Blue lines indicate nuclear segmentation result. Scale bars: 20 µm in the left image, 2 µm in the right zoom-in image. B, Correlation between RAEFISH and scRNA-seq results. **C-E,** Spatial distribution of marginal zone B cells (**C**), follicular B cells (**D**) and dendritic cells (**E**).

## Methods

### Probe design and synthesis

Each of our RAEFISH probe libraries for profiling endogenous human and mouse transcriptomes contained three sub-libraries: a reverse padlock probe sub-library, a linear priming probe sub-library, and an encoding probe sub-library. Each reverse padlock probe contained from the 5’ to 3’: a 20-nt common region, a 30-nt RNA-binding region (reverse-complement of the target RNA sequence), and another 30-nt common region. The total length of the reverse padlock probe is 80 nt. Each linear priming probe contained from the 5’ to 3’: a 20-nt common region, a 30-nt RNA-binding region, and a 10-nt common region that binds to the reverse padlock probe to prime rolling circle amplification. The total length of each linear priming probe used in the RCA step is thus 60 nt. To allow cost-efficient pooled synthesis of these linear priming probes, we further added to their 3’ ends a 47-nt common sequence containing the reverse complement of a T7 promoter (27 nt) and a PCR priming region (20 nt). So in total, the length of each probe in our un-amplified linear priming probe sub-library is 107 nt. Each encoding probe contained from the 5’ to 3’: a 20-nt common region, a 20-nt readout probe binding region, a second 20-nt readout probe binding region, a 30-nt sequence same as the RNA-binding region on the corresponding reverse padlock probe, a third 20-nt readout probe binding region, a fourth 20-nt readout probe binding region, and another 20-nt common region. The total length of each encoding probe is 150 nt. Each gene was encoded with a unique 94-bit Hamming Distance 4 (MHD4) binary code with Hamming Weight 4 – having four “1” bits corresponding to the four readout probe binding sequences on the encoding probe. Because our probe amplification procedure reverse-complements this sequence, the un-amplified encoding probe sequences in the encoding probe sub-library were reverse-complemented. We require that on the target RNA the reverse padlock probe and the linear priming probe bind right next to each other, without overlap in their binding sequences and with no or a 1-nt gap. We were able to design probes targeting 23,312 human genes, including 16,501 protein-coding genes and 6,811 long-noncoding RNAs from Gencode v46, each probed by one set of reverse padlock, linear priming, and encoding probes. We were able to design probes targeting 21,955 genes in the mouse transcriptome, including 16,618 protein coding genes and 5,337 lncRNAs from Gencode vM35. Most mouse genes were probed with two sets of reverse padlock, linear priming, and encoding probes. For the minority of the mouse genes that only yielded one pair of good binding regions for the reverse padlock and linear priming probes, we duplicated the reverse padlock, linear priming, and encoding probes in the sub-libraries as a compensation. The un-amplified reverse padlock, linear priming, and encoding probe sequences for human and mouse transcriptome RAEFISH were ordered as oligo pools from Twist Bioscience. 94 readout probes with adapter sequences that bind two different dye-labeled probes, as well as dye-labeled probes themselves, were ordered from Integrated DNA Technologies (IDT). The sequences of the ordered probes are listed in Table S3.

For the Perturb-RAEFISH probe design, the reverse padlock probes contained 40-nt RNA-binding regions, created by combining the 20-nt gRNA spacer sequences with a 20-nt constant sequence from the upstream U6 promoter. The shared linear priming probe binds directly to the downstream gRNA scaffold with a 45-nt RNA-binding region. To design the encoding probes, we assigned a unique 14-bit Hamming Distance 2 (HD2) binary code with Hamming Weight 4 to three gRNAs for the same perturbation target gene. The encoding probes contain 40-nt rolling circle amplicon binding regions, corresponding to the 40-nt RNA-binding regions on the reverse padlock probes. The rest of reverse padlock probes, linear priming probe and encoding probes were assembled similarly as in endogenous RNA RAEFISH probe assembly. The un-amplified reverse padlock and encoding probe sequences for gRNA RAEFISH were ordered as oligo pools from Twist Bioscience (Table S11). The single shared linear priming probe, the 14 dye-labeled readout probes were ordered from IDT (Table S11).

For the MERFISH probe design, we selected 492 target genes, with a broad range the FPKM in A549 cells and included 183 genes that were targeted by gRNAs in CRISPR screen. Next, the MERFISH probe library was designed following the procedure as described in previous publications^15,111,112^. Each gene was assigned a unique 26-bit MHD4 code. The MERFISH encoding probe template library was ordered as an oligo pool from Twist Bioscience (Table S12). The 26 readout probes with adapters and two dye-labeled probes that bind to the adapters were ordered from IDT (Table S12).

To amplify the oligo pools, we used a previously reported experimental workflow^111^. Briefly, this workflow involves limited-cycle PCR, in vitro transcription, reverse transcription, alkaline hydrolysis and oligo purification. Specifically, we used a phosphate-labeled primer for reverse transcription during padlock probe synthesis, to enable the circularization step with the T4 ligase in RAEFISH sample preparation. Limited-cycle PCR primers, and reverse transcription primers were ordered from IDT. The sequences are detailed in Table S13.

### RAEFISH protocol for A549 endogenous transcriptome

A549 cells (ATCC, CCL-185) were cultured on coverslips (Bioptech; 40-1313-0319) for 3– 4 days and subsequently fixed with 4 mL of 4% paraformaldehyde (PFA) at room temperature (22°C) for 20 minutes (min). The 4% PFA was prepared by diluting 16% PFA solution (Electron Microscopy Sciences; 15710) with Dulbecco’s Phosphate Buffered Saline (DPBS, Sigma-Aldrich; D8537-500ML). After fixation, the samples were washed twice with 3 mL of DPBS for 2 min each. The samples were then permeabilized with 3 mL of 0.5% Triton X-100 solution (Sigma-Aldrich; 93443) in DPBS for 10 min. After permeabilization, the samples were washed three times with 3 mL of DPBS, for 3 min each time. Then, they were washed twice with 3 mL of 2×SSC, prepared by diluting 20×SSC buffer (Invitrogen; 15557-044) with double-distilled water (ddH_2_O), for 2 min each time. For the prehybridization, the samples were incubated with 3 mL prehybridization buffer (in 2×SSC, 50% formamide (Sigma-Aldrich; F7503-1L), 0.1% Tween 20 (Sigma; P7949-500ML), 120 units of Murine RNase Inhibitor (MRI, New England Biolabs; M0314L)) at room temperature for 10 min. After prehybridization, each sample was placed face-down and incubated with 50 µL primary hybridization buffer (2×SSC, 50% formamide, 20 U MRI, 3 nM per padlock probe and priming probe) in a 37°C incubator for 16 hours. Then, the samples were washed with 2 mL of 50% formamide (in 2×SSC, 0.1% Tween 20 and 120 U MRI) three times at 37°C, for 15 min each. Subsequently, samples were washed with 3 mL DPBS twice at room temperature, for 2 min each.

Next, samples were placed face-down and incubated with 100 µL T4 DNA ligation mixture (50U T4 ligase (Thermo Scientific; EL0011), 1 µM splint oligo, 1 mM DTT, 1 mM ATP (Thermo Scientific; R0441), 0.2 mg/mL recombinant albumin (BSA, New England Biolabs; B9200S), 40 U MRI, and 1×T4 DNA ligase buffer in ddH_2_O) at room temperature for 2 hours. The 16-nt splint oligo (GAGAGTGGGTATA*T*C/3InvdT/) was designed to bind two 6-nt sequences on both ends of the padlock probe to enable padlock probe circulation, and also have a 3-nt overhanging sequence with phosphorothioate modification and an inverted dT at its 3’ end to prevent it from unwantedly functioning as a primer for RCA. After ligation, the samples were first washed with 2 mL of 20% formamide (in 2×SSC buffer) in a 37°C water bath for 15 min, followed by two times of 2 mL of toehold washing buffer (20% formamide, 2×SSC and 800 nM toehold oligo (GA+TAT+ACC+CAC+TC+TC, “+” represents locked nucleic acid)) in a 37°C water bath, for 15 min each time to further remove extra splint oligos. The samples were subsequently washed three times with 3 mL of DPBS at room temperature for 5 min each. Then, the samples were placed face-down and incubated with 50 µL RCA mixture (50U Phi29 DNA Polymerase (Thermo Fisher; EP0092), 0.5mM Deoxynucleotide (dNTP) Solution Mix (New England Biolabs; N0447S), 1mM DTT, 0.2 mg/ml BSA, 20 units MRI and 1×Phi29 buffer) at 30°C for 2 hours. After the RCA step, the samples were briefly washed with 3 mL DPBS at room temperature, and fixed again with 3 mL 4% PFA in PBS for 30 min at room temperature. Then, the samples were quickly washed once with 3 mL DPBS at temperature. Next, each sample was incubated with 50 µL of hybridization buffer (2×SSC, 40% formamide, 10% dextran sulfate (Sigma, D8906-50G), 20 U MRI and 0.1 mg/mL yeast tRNA (Invitrogen, AM7119)) with 3 nM per probe of the human RAEFISH encoding probes at 37°C for 16 hours. After the encoding probe hybridization, the samples were washed twice with 2 mL of 40% formamide (in 2× SSC and 0.1% Tween) in a 37°C water bath, for 15 min each time. This was followed by a single wash with 3 mL of 40% formamide (in 2× SSC) for 2 min at room temperature. To account for sample drift during sequential hybridization and readout imaging, 0.1-µm yellow-green fiducial beads (Invitrogen, F8803) were incubated on the surface of the samples. Finally, each sample was mounted in Bioptech’s FCS2 flow chamber to enable automatic buffer exchange and sequential imaging as described before^112^.

### RAEFISH protocol for mouse tissue

The C57BL/6 mice lineage were rapidly anesthetized with isoflurane and subsequently euthanized. The liver and lymph nodes were collected, promptly embedded in O.C.T. compound, and rapidly frozen using dry ice before being stored at -80°C for long-term preservation. Pregnant mice were euthanized on embryonic day 13.5 (E13.5) using the same euthanasia and tissue embedding method to collect placentas. For the tissue sectioning, the tissue blocks were equilibrated in a cryostat (Leica CM3050S) at -20°C for 20 min before sectioning. The liver and placenta were sectioned at 10 µm, while lymph nodes were sectioned at 8 µm at -20°C. These slices were attached on 40-mm coverslips pretreated with a 1:1 mixture of 34–37% (v/v) hydrochloric acid (Sigma-Aldrich; HX0607-1) and methanol (Sigma-Aldrich; 179337-4L-PB), followed by poly-D-lysine solution (Millipore, A-005-C) treatment at room temperature (22°C).

The tissue slices from all three types were fixed with 4 mL of 4% PFA in PBS at room temperature for 30 min. After fixation, the slices were washed three times with 3 mL of 2×SSC buffer, prepared by diluting 20×SSC buffer with ddH_2_O, with each wash lasting 3 min. The slices were then permeabilized with 3 mL of 0.5% Triton X-100 solution with DPBS for 20 min at room temperature. Following permeabilization, the slices were washed three more times with 2×SSC buffer, 3 min for each time. Prehybridization was performed using 3 mL of 50% formamide in 2×SSC buffer with 0.1% Tween 20, and 60 U of MRI. For hybridization, the slices were placed face-up, and each slice was incubated with 25 µL of hybridization buffer (in 2×SSC, 50% formamide, 10% dextran, 0.1 mg/mL yeast tRNA, 10 U of MRI, pooled reverse padlock and linear priming probes at 3 nM per oligo), and covered with a piece of parafilm in a 37°C incubator for 16 hours. Then, the slices were washed three times with 2 mL of 50% formamide (with 400 units MRI and 0.1% Tween 20) at 37°C in a water bath, 15 min for each wash. The slices were then washed once with DPBS for 3 min and briefly rinsed with DPBS at room temperature.

Next, the slices were placed face up and incubated with 100 µL of Hi-T4 DNA ligation mixture (containing 4000 U of Hi-T4™ DNA Ligase (New England Biolabs; M2622L), 1 µM splint oligo, 1 mM DTT, 1 mM ATP, 0.2 mg/mL BSA, 40 U of MRI, and 1× Hi-T4 DNA ligase buffer) at room temperature for 16 hours, with parafilm covering to prevent evaporation. After ligation, the slices underwent the same toehold washing procedure and utilized the same toehold oligo described in the “RAEFISH protocol for A549 endogenous transcriptome.” Then, the slices were placed face-up and incubated with 100 µL RCA mixture (100 U Phi29 DNA polymerase, 0.5 mM dNTP, 1 mM DTT, 0.2 mg/ml BSA, 40 U of MRI and 1×phi29 buffer) at 30°C for 3 hours. After the RCA step, the slices were quickly washed with 3 ml DPBS at room temperature, and fixed again with 3 mL 4% PFA in PBS for 30 min at room temperature. Then, the slices were quickly washed with DPBS and incubated with encoding probe prehybridization buffer (40% formamide in 2×SSC) at room temperature for 2 min. After encoding probe prehybridization, the slices were incubated with 25 µL encoding hybridization buffer (2×SSC, 40% formamide, 10% dextran sulfate, 400 units MRI and 0.1 mg/mL yeast tRNA, pooled encoding probes at 3 nM per oligo) at 37°C for 16 hours. After the encoding probe hybridization, the remaining steps were then performed as described in the “RAEFISH protocol for A549 endogenous transcriptome”.

### CRISPR screen library construction

We selected 574 DNA/chromatin binding nuclear protein coding genes that were expressed in A549 and designed 3 gRNAs for each target. The gRNA spacer sequences with PCR priming regions on the 5’ and 3’ ends were order in an oligo pool from Twist Bioscience (Table S9). To construct the plasmid library, PCR amplified gRNA library were assembled with a FastDigest Esp3I (Thermo Fisher Scientific, FD0454) digested backbone through Gibson Assembly. The backbone was modified from plasmid CROPseq-Guide-Puro^101^ (Addgene, 86708), and its sequence is included in File S1. Lentivirus was produced in HEK-293FT cells (Thermo Fisher Scientific, R70007) and was used to transduce an engineered A549 clone with dox-inducible Cas9 (Addgene, 83481) expression. Cells were under Puromycin (Thermo Fisher Scientific, A1113803) selection for one week until the cell library was established, and were always maintained in medium with Puromycin. To perform CRISPR knockout screen, Cas9 expression was induced with 1 µg/mL doxycycline (Dox) for one week before cells were seeded onto coverslip for experiments.

### Perturb-RAEFISH protocol in A549 cells

The screen library A549 cells with induced Cas9 expression were seeded on coverslip for 3-4 days and fixed with 4% PFA in DPBS at room temperature for 20 min. After fixation, the samples were washed twice with DPBS for 2 min each. The samples were then permeabilized with a 0.5% Triton X-100 solution in DPBS for 10 min. After permeabilization, the samples were washed three times with DPBS, for 3 min each time. Then, they were washed twice with 2×SSC for 2 min each. For the prehybridization, the samples were incubated with 3 mL prehybridization buffer (30% formamide, 0.1% Tween 20, 120 U of MRI) at room temperature for 10 min. After prehybridization, each sample was placed face-down and incubated with 50 µL primary hybridization buffer (2×SSC, 30% formamide, 200 U of MRI, 5 nM per padlock probe and 100 nM priming probe) at 37°C for 16 hours. Then, these samples were washed with 3 mL of 45% formamide (in 2×SSC, 0.1% Tween 20 and 120 U of MRI) three times, for 15 min each. Subsequently, samples were washed with DPBS twice, for 2 min each. Next, samples were placed face-down and incubated with 50 µL T4 DNA ligation mixture (2000 U of Hi-T4 ligase, 1 µM splint oligo, 1 mM DTT, 1 mM ATP, 0.2 mg/mL BSA, 20 U of MRI, and 1× Hi-T4 DNA ligase buffer) at room temperature for 2 hours. Afterward, the same procedures for toehold washing, RCA, and encoding probe prehybridization were followed as outlined in the “RAEFISH protocol for A549 endogenous transcriptome”. To enable the RNA MERFISH readout in the same sample, each sample was incubated with 50 µL of hybridization buffer (2×SSC, 40% formamide, 10% dextran sulfate) with both 3 nM per probe of the RAEFISH encoding probes and 0.5 nM per probe of the MERFISH encoding probes at 37°C for 16 hours. The remaining steps were then performed as described in the “RAEFISH protocol for A549 endogenous transcriptome”.

### Sequential secondary probe hybridization and imaging

For the whole-genome-scale transcriptome RAEFISH readout (94 bits), we performed 47 rounds of readout probe hybridization and dual-color imaging; For Perturb-RAEFISH readout (14 bits), 7 rounds of dual-color imaging were formed; For RNA MERFISH readout (26 bits), 13 rounds of dual-color imaging were formed. Sequential hybridization and imaging were perform as described in previous studies^101,112^. RNA MERFISH imaging was performed before Perturb-RAEFISH imaging. Both RAEFISH and RNA MERFISH signals were captured using the 750-nm and 647-nm laser channels, while the yellow-green fiducial bead signals were collected using the 488-nm laser channel. Each imaging z stack covered a total z-depth of 12 μm. To remove and block signals from the previous round of imaging, the samples were washed with 65% formamide in 2× SSC with 1 mM blocking oligos (Table S3) that were complementary to the dye-labeled readout-adapter-binding oligos.

### Data analysis

#### Decoding of endogenous RAEFISH and MERFISH signals

We adopted a previously reported pixel based decoding strategy (PMID: 27625426). First, the drifts between RAEFISH hybridization rounds were corrected by 2D fitting of fiducial beads. Next, for each z-stack image, the background was removed by subtracting an image generated by image opening with a disk-shaped morphological structuring element with a radius of four or five pixels. The values of each pixel in 94 rounds of imaging were concatenated into a 96-dimensional “pixel vector”. The pixel vectors were normalized to generate unit vectors and compared with unit vectors of all 25,281 possible 94-bit MHD4 codes. The closest possible code for each pixel was identified if it is within a code vector distance threshold of 0.65. The adjacent pixels matched to the same code were combined as one molecule. To further select real RNA molecules, we created a parameter space including three properties: the foci area, the distance to the matched code, the magnitude of the pixel vector before normalization. The parameter space was segmented into small bins and the error rate of each bin was calculated by the ratio of foci number that matched to empty control codes over foci number that matched to all 25,281 possible codes. The foci with parameters in bins with an error rate smaller than a cutoff threshold will be finally selected. For each dataset, 8 random field of views were used to define the parameter space. Finally, all the RAEFISH foci within the selected parameter bins were decoded. RNA MERFISH was decoded following the same strategy as above.

#### Decoding of Perturb-RAEFISH signals

First, drift correction was perform as mentioned above. Background removed RAEFISH signals from 14 rounds were projected onto a 2D images and the bright RAEFISH foci were segmented. Next, segmented RAEFISH foci were matched with 1,001 possible 14-bit HD2 codes. Only foci with a top-4 normalized “pixel vector” element exceeding 0.1 and a top-6 element below 0.1 were selected.

#### Cell segmentation

For mouse liver, we perform cell segmentation with a combination of CellPose^36-38^ and ClusterMap^39^. For mouse placenta and lymph node, we performed cell segmentation with CellPose. For Perturb-RAEFISH, cells were segmented with a watershed algorithm.

#### Zonation score analysis

For liver zonation analysis, a 50-μm radius was defined around each cell, and the zonation score was calculated as the percentage of nearby periportal hepatocytes among all hepatocytes within this radius. The cells without any hepatocytes in the radius was assigned a zonation value of 0.5. For lymph node zonation analysis, a 150-μm radius was defined around each cell, and the zonation score was calculated as the percentage of nearby B cells (including marginal zone B cells, follicular B cells and germinal center B cells) among both B cells (the same above) and T cells (T cell major group) within this radius.

#### Cell-cell interaction analysis

To analyze cell-cell interaction enrichment, an interaction distance threshold of 20 μm was used, and each pair of cells within this range was considered interacting. We calculated the interaction enrichment between each pair of cell types by the following equation:

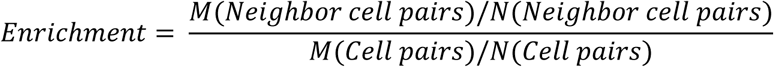

*M*(*Neighbor cell pairs*) represents the number of observed neighboring cell pairs of the two cell types, *N*(*Neighbor cell pairs*) represents the number of all observed neighboring cell pairs in the whole tissue regardless cell types, *M*(*Cell pairs*) represents the number of all possible cell pairs between the two cell types given their cell numbers, and *N*(*Cell pairs*) represents the number of all possible cell pairs in the whole tissue given the total cell number.

## Supplementary Materials

Table S1. Human endogenous transcriptome RAEFISH codebook and template probe libraires, related to Fig. 1 and 2.

Table S2. A549 cell-cycle associated genes, related to Fig. 2.

Table S3. Mouse endogenous transcriptome RAEFISH codebook, template probe libraries, and the corresponding readout and blocking oligos (used for both human and mouse endogenous transcriptome RAEFISH), related to Fig. 1-6, S1-3.

Table S4. Liver zonation score correlated genes in each cell type, related to Fig. 3.

Table S5. Differentially expressed genes (DEGs) of cholangiocytes neighboring versus not neighboring leukocytes, related to Fig. 4.

Table S6. Differentially expressed genes (DEGs) of leukocytes neighboring versus not neighboring cholangiocytes, related to Fig. 4.

Table S7. Differentially expressed genes (DEGs) of macrophages neighboring versus not neighboring decidual stromal cells, related to Fig. 5.

Table S8. Differentially expressed genes (DEGs) of decidual stromal cells neighboring versus not neighboring vascular endothelial cells, related to Fig. 5.

Table S9. gRNA template oligo library for Perturb-RAEFISH lentiviral plasmid library construction, related to Fig. 7.

Table S10. Hierarchically clustered perturbation target gene and MERFISH-probed gene names, related to Fig. 7. Cluster IDs are shown. A cluster ID of -1 indicates the gene is not in a cluster.

Table S11. Perturb-RAEFISH codebook, template probe libraries and readout oligos, related to Fig. 7.

Table S12. RNA MERFISH codebook, template probe library, and the corresponding readout and blocking oligos, related to Fig. 7.

Table S13. Oligos used for library amplification, related to all figures.

File S1. Plasmid backbone sequence of CRISPR screen library, related to Fig. 7.

## Notes

### Summary of Updates

Some typographical errors have been corrected in this version.

